# Catcher in the rye: museomics reveals the identity of cereal remains recovered from a late nineteenth century antique furniture

**DOI:** 10.64898/2026.09.04.749326

**Authors:** Péter Poczai, Saeideh Javid, Zoárd Kenessey, Sundre Winslow, Laura Capelatti, Rocio Deanna, Tamás Németh, Otto Veisz, Ildikó Karsai, András Cseh

## Abstract

Historical crop remains preserved in everyday objects are an underused archive of agrobiodiversity change, yet their degraded DNA resists conventional analysis. We recovered cereal grains and straws used as upholstery padding in a Central European settee dated to about 1870 to 1900. We identified and characterized the specimen by genome skimming. Damage-aware sequencing confirmed authentic historical DNA of modest, late nineteenth-century age. The assembled plastome placed the specimen securely within rye, and a genome-wide SNP panel of rye landraces resolved it inside cultivated *Secale cereale* subsp. *cereale*, closest to a Hungarian bred rye and a Slovak upland landrace and remote from wild or eastern lineages. Screening 192 stress-associated loci recovered the structural machinery of resistance, abiotic stress and aluminum tolerance pathways, while heading-date genotyping showed the Weining earliness allele at Hd2R already present. Divergence across 36 domestication sweep regions concentrated at a few improvement loci. Integrating these genomic signals with genebank passport data and an 1891 newspaper record, we interpret the specimen as a tall Central European rye at the threshold between landrace and improved variety. Historical crop material, read this way, preserves both genomes and the traces of human agricultural practice.

## Introduction

Museomics, broadly understood as the molecular investigation of historical collections, has emerged as a transformative discipline by enabling the extraction, sequencing, and analysis of degraded historical DNA (hDNA) from specimens preserved in musea or repositories [1–5]. While ancient DNA (aDNA) typically refers to genomic material from long-deceased organisms preserved under permafrost or archaeological contexts (>500 years), hDNA refers to more recently degraded DNA, often less than 500 years old, extracted from specimens preserved in musea or herbarium collections under controlled or archival conditions. Museomic approaches thus extend the traditional domain of ancient DNA (aDNA) by encompassing more recent, yet still fragmented, genomic material, offering direct insight into past biological diversity, evolutionary trajectories, and responses to environmental change [3,6–8]. The advent of high-throughput sequencing technologies, notably genome skimming, circumvents limitations of PCR-based barcoding by enabling short-read sequencing of total DNA from degraded specimens, generating large quantities of genomic data suitable for assembly of chloroplast genomes, nuclear ribosomal DNA, and extended barcode loci [9–11]. Compared to the conventional single-barcode strategy, genome skimming is especially advantageous for museomic applications because it avoids PCR failures due to primer mismatches or DNA fragmentation, and can provide resolution often below the species level [12,13].

Cereals, including major crops and their wild relatives, form an integral part of our global food security framework. Among these, landraces, locally adapted traditional varieties, represent reservoirs of genetic diversity that are critical for crop improvement, resilience to biotic and abiotic stress, and adaptation to changing environmental conditions [14,15]. Historical cereal specimens preserved in musea and herbaria thus offer a unique opportunity to explore genetic diversity through time, encompassing lost alleles, extinct lineages, and shifts due to domestication or breeding [6,16]. By applying museomic approaches to cereal landraces and wild relatives, one can trace the temporal dynamics of genetic variation, assess the maintenance or erosion of diversity, and inform both evolutionary understanding and practical conservation or breeding efforts [17,18].

Previous studies that deployed historical DNA analyses in cereals have demonstrated the feasibility of recovering genomic data from museum specimens, enabling retrospective comparisons with modern varieties, reconstruction of past populations, and elucidation of evolutionary processes underpinning domestication and adaptation [19–24]. The ability to extract informative genetic markers and organellar genomes from century-old cereal-remains reinforces the utility of museum collections as archives for agricultural and evolutionary investigations. The synergy of museomics and genome skimming potentiates the generation of extensive reference datasets for cereals that are crucial for taxonomic verification, genotype to phenotype mapping, and conservation of genetic resources.

Here we apply a museomic, genome-skimming based approach to rye straws recovered from the upholstery padding of a late nineteenth century antique settee. Combining morphological examination with historical DNA sequencing, we recover and assemble a high-coverage, complete plastid genome from this degraded material and confirm its fidelity by independently amplifying the inverted repeat junctions and indel regions. We then identify the specimen through its morphology, the architecture and phylogenetic position of its plastome, and genome-wide nuclear variation that places it among rye landraces and points to its likely origin. In doing so, we show that plant material repurposed within historical objects offers a still underused archival source for the museomics of cereals.

## Materials and methods

### DNA extraction

Historical plant tissue yields short, damaged and easily contaminated DNA, so all work on the old cereal specimen was carried out in facilities dedicated to historical DNA. Small leaf segments of 2 to 4 mm^2^ were removed from the recovered historical cereal specimen (Rozs3) in 2018 and extracted in the Historical DNA Laboratory of the Finnish Museum of Natural History. Leaf samples were placed in 2.0 mL Eppendorf Safe-Lock tubes with laboratory grade quartz sand (Merck, Darmstadt, Germany) and two sterile steel beads, then homogenized to a fine powder in a TissueLyser II (Qiagen, Venlo, The Netherlands) at room temperature. Extraction followed the CTAB protocol of Doyle and Doyle [25], and two further extractions used the DNeasy Plant Mini Kit (Qiagen, Venlo, The Netherlands) with the homogenization described above. As a fresh reference, seeds of the rye variety Lovászpatonai [26] were sown in experimental pots and raised in the greenhouse of the Kaisaniemi Botanical Garden, and seedling DNA was extracted with the same DNeasy protocol. DNA quality and quantity were assessed by NanoDrop microvolume spectrophotometry (Thermo Fisher Scientific, Waltham, MA, USA) and a Qubit 4 fluorometer (Thermo Fisher Scientific, Waltham, MA, USA), and DNA integrity was measured as the DNA integrity number (DIN) on a TapeStation 4150 (Agilent Technologies, Santa Clara, CA, USA). The cereal remains are deposited in the collection of the HUN-REN Centre for Agricultural Research (HUN-REN ATK), Martonvásár, Hungary.

### Library construction and sequencing

For the historical extractions, four separate TruSeq DNA Nano libraries (Illumina, San Diego, CA, USA) were built by Novogene (Munich, Germany) following the manufacturer’s protocol. Because the template was already short, the mechanical fragmentation step was omitted and the intact fragments were carried straight into end repair, A tailing and adapter ligation, followed by indexing amplification and a bead-based clean up. Extracted DNA was submitted at about 90 ng per sample, as 50 µL at 1.8 ng/µL. Of the four libraries two passed quality control and were retained, and these were sequenced on a NovaSeq X Plus System with 2 x 150 bp paired end reads at the same facility. The freshly extracted Lovászpatonai DNA was prepared and sequenced by the same protocol. We deliberately applied no enzymatic damage repair, for example, the partial uracil-DNA-glycosylase treatment of Rohland et al. [27], because herbarium DNA degradation kinetics remain little studied [28] and we wished to preserve the raw damage signal for such future work. Reads deposited under BioProject PRJNA1510302.

### Quality control and read processing

Historical libraries carry laboratory and environmental DNA alongside the target genome, so reads were screened for contaminants before any downstream analysis. Raw reads were passed through FastQ Screen with its default database [29], and a custom database was built from the laboratory contaminants and post-mortem colonizing microorganisms reported by Glassing et al. [30] and Bieker et al. [31] to remove exogenous DNA according to Winslow et al. [13]. The corresponding genomes were downloaded from NCBI [32], indexed with bowtie2 [33] and added to the FastQ Screen configuration, and low complexity reads matching multiple genomes were discarded. Low-quality bases and adapters were then removed with BBDuk from the BBTools package [34] at a minimum quality of 20 and a minimum length of 60 bp, with read quality reassessed in FastQC [35].

### Chloroplast genome assembly and analyses

The maternally inherited plastome behaves as a single linked marker for placing the specimen, so we assembled it first. However, discordance has been recorded among plastid genes [36]. Paired end Illumina reads were assembled de novo with GetOrganelle v1.7.5 [37] using the embplant_pt seed database (-F) over 15 extension runs (-R) and default SPAdes kmer settings (-k 21,45,65,85,105) [38], and assembly graphs were inspected in Bandage [39]. To verify the indels detected in the assembly, we designed primers in Geneious Prime against the Lovászpatonai rye reference, including four primers amplifying a 1500 bp and a 550 bp fragment across the *rbcL-accD* and *rpl2-ycf2* intergenic spacers. PCR contained 20 ng of template DNA, 2 X GoTaq Master Mix (Promega, USA) and 0.2 uM of each primer in a total volume of 25 ul. Amplification was carried out in an Eppendorf Mastercycler (Eppendorf, Germany) using the following profile: 90 s at 94°C; 36 cycles of 30 s at 94°C, 40 s at 60°C, 40 s at 72°C and a final extension at 72°C for 3 min [40]. Amplicons were separated on 1.5% agarose gels in 0.5 x TBE buffer (220 V, 0.5 h) and stained with ethidium bromide. Sanger sequencing used an ABI 3730XL sequencer with the ABI PRISM BigDye Terminator kit. Genomes were annotated with GeSeq [41], visualized with Chloroplot [42] and their inverted repeat junctions plotted with IRplus [43], then deposited in NCBI under accession numbers PZ895511-PZ895512. For phylogenetic placement we used the plastid genomes of wheat (NC_002762) [44], barley (NC_008590), sorghum (NC_008602) [45], oat [46], rice (NC_031333) [47], maize (NC_001666) [48], and millet (NC_029732), with pineapple (NC_026220) [49] as an outgroup. Because grasses and bromeliads carry three major plastome inversions [50], which we plotted with shinyCircos-V2.0 [51], we extracted 70 coding regions and aligned them with MAFFT v7.49 [52]. Maximum likelihood analysis was run in RAxML v8.2.11 [53] under GTR+Γ with 1,000 bootstrap replicates, treating the plastid genes as a single partition representing one heritable unit.

### Genome wide SNP genotyping and phylogenetic placement

To locate the origin of the recovered specimen we placed it within a genotyping by sequencing (GBS) panel of rye landraces. We downloaded the GBS datasets of Schreiber et al. [54] and Rabanus-Wallace et al. [55] covering landraces from the Genebank of the Leibniz Institute (IPK), Gatersleben, Germany (Supplementary Table S1), concentrating on the regional spread of landraces and varieties, and excluding other *Secale* taxa apart from *S. strictum* and S*. vavilovii*, which served as outgroups. Reads were demultiplexed and quality filtered with the process_radtags module of STACKS v2.68 [56], and a genome wide nuclear matrix was assembled following the phylogenomic protocol of Rick et al. [57].

All pre-processed reads were mapped to the chromosome scale reference of Weining rye [58] with bowtie2 in fast and memory efficient mode (--very-fast-local), and the alignments were converted and sorted with samtools [59]. Historical DNA accumulates atypical nucleotides, interstrand cross links, and strand breaks through oxidative and hydrolytic decay [60–62], and these lesions can mimic true variants. To keep such post-mortem misincorporations from being miscalled as SNPs, we estimated the mean overhang length (λ), nick frequency (ν) and cytosine deamination rates in double stranded regions (δd) and overhangs (Cend) for the specimen under the Bayesian framework of mapDamage2 [63], then rescaled the base quality scores in the BAM files by their probability of arising from damage.

Single nucleotide polymorphisms were written in variant call format (VCF) [64] with mpileup v1.6 [65], and variants were filtered and called in bcftools v1.9 [66], retaining sites with a genotype quality above 10 and a mapping quality above 40. Genotypes were called only at a read depth between 5 and 250 with a minor allele count above 3, and no missing data were allowed in the nuclear matrix. The filtered VCF was converted to PHYLIP with vcf2phylip v2.0 [67], invariant sites were removed with the raxml_ascbias.py script (https://github.com/btmartin721/raxml_ascbias), and maximum likelihood analysis was run in IQ-TREE v2.3.6 [68] under an ascertainment bias corrected model (GTR+ASC) [69]. The best tree was taken from 10 runs of 1,000 ultrafast bootstrap replicates [70] with SH like approximate likelihood ratio tests [71], and a hill climbing nearest neighbor interchange search (-bnni) curbed overestimation of support. Trees were displayed and edited in the Interactive Tree of Life v6 (iTOL) [72]. All analyses were run on the Puhti supercomputer at CSC, Espoo, Finland. The SNP matrix (Supplementary Material 2) also supplied a principal component analysis using the *adegenet* package [73,74] in R [75].

### Screening for stress associated loci of rye

To assess the trait content of the historical rye specimen we screened public databases for characterized stress associated loci of rye. Working from the primary literature, we retrieved NCBI records for genes and markers implicated in biotic stress resistance, in abiotic stress response, and in aluminum tolerance, and assembled them into a curated reference set. The set spanned nematode, aphid, rust, and powdery mildew resistance genes, as well as resistance gene analogues, herbicide target loci, rye stress responsive transcripts, patent derived stress tolerance candidates. The aluminum tolerance system at the Alt3, Alt4, and Qalt5 loci, including the ALMT1 malate transporter cluster and the ScAACT1 (ScMATE) citrate transporter (Supplementary Material 3). Quality filtered reads from the specimen were mapped to each reference with the Geneious assembler (default settings), and a consensus was called per locus. For every reference we recorded whether reads mapped, the recovered consensus length and its gap count, the lengths of the individual recovered blocks, the number of substitutions in the consensus relative to the reference, and the maximum per site read depth. A locus was scored as recovered when at least one read was mapped and a consensus formed, and partial recoveries were kept. Recoveries returned as two or more separate blocks were flagged as fragmented, since block structure tracks the short insert sizes expected from degraded template. The annotated reference set and per locus metrics are given in Supplementary Material 3. Summary figures were produced in Python with matplotlib [76], pandas [77], and NumPy [78].

### Heading date QTL genotyping

Because flowering time separates old landraces from modern cereals, we asked whether the specimen already carried the early heading alleles of rye. Li et al. [58] mapped three heading date QTLs in a Weining by Jingzhou cross, Hd2R on chromosome 2R over the photoperiod gene ScPpd1, Hd5R on 5R and Hd6R on 6R, whose Weining alleles act additively towards earliness. We extracted the genomic window around the diagnostic site of each QTL from the Weining assembly and mapped the historical specimen reads to the three windows in Geneious Prime (Biomatters Ltd, Auckland, New Zealand). At each diagnostic position, we recorded the read depth, the base composition and its split across forward and reverse strands, and read the specimen allele against the Weining earliness state.

### Variation at domestication associated sweep regions

To ask whether the historical rye departs from modern rye at loci shaped by domestication, we examined the selective sweep regions defined in the Weining genome by Li et al. [58] from the Derived Ratio Index (DRI), the cross-population statistic (XP) and FST-based differentiation. We extracted the 36 sweep regions from the Weining assembly, mapped the Rozs3 reads to them and called variants to count the polymorphic sites separating the historical rye from the reference. We then built a reference guided consensus of the historical rye for each region and used it as the target for mapping whole genome resequencing reads of three modern cultivars, R2446 (ERR3771531), R925 (ERR3771534), and PUMA (SRR10088797), from Rabanus-Wallace et al. [55]. SRA files were converted to FASTQ with fastq-dump from the SRA Toolkit, and variants were called for each cultivar against the historical consensus with the SNP caller of Geneious Prime (Biomatters Ltd, Auckland, New Zealand) under default settings, so that every count express polymorphism relative to the historical rye.

## Results

### Origin and morphological identification of the recovered cereal specimen

The cereal was recovered from an antique settee assessed by an art historian and an antiquarian dealer as a *canapé* in the Habsburg Biedermeier manner, more precisely a later Biedermeier revival of the Historicist or Gründerzeit period dating from about 1870 to 1900 (Figure S1). The piece has since been restored and reupholstered (Figure S1B), but at the time of sampling its original upholstery layers were undisturbed (Figure S1A). Both rye ears and loose straw had been packed as seat and back padding beneath an intact hessian and linen cover, a sealed context that minimized modern contamination and supports the historical integrity of the material.

Morphologically, the packing material was rye (Figure 1). The recovered ears (Figure 1C) were long, slender and somewhat lax, tapering toward the apex and bearing long flexuous awns, unlike the denser, shorter spike of the modern rye reference (Figure 1A) or the broader, more compact awned wheat (Figure 1B). The straw was long and fine with elongated internodes, consistent with the tall growth habit of older landrace rye. The caryopses (Figure 1F) were narrow and elongate with a pointed apex, an apical hair brush, and a deep ventral groove typical of *Secale*, and were clearly slenderer than the plump, rounded grains of wheat (Figure 1E). In spike, straw, and grain the specimen matched an old, tall, awned rye landrace.

### Authenticity and post-mortem damage of the historical DNA

To confirm the authenticity of the recovered sequences and characterize their preservation, we screened the historical reads for the damage signatures of degraded DNA with mapDamage2.0 [63]. Fragment lengths were short, with read occurrence highest at the smallest sizes and declining steadily toward longer molecules across the 26 to 176 bp range examined, as expected for degraded historical material (Figure S2). The misincorporation profile carried the diagnostic pattern of ancient and historical DNA, a detectable excess of C to T substitutions toward the 5′ ends of reads and of complementary G to A substitutions toward the 3′ ends, indicating genuine deaminated template rather than modern contamination (Figure S3). The magnitude of this damage was nonetheless low. The Bayesian damage model converged well over 50,000 iterations (Figures S4 to S6) and estimated a single stranded overhang parameter λ of about 0.16 together with very low cytosine deamination rates (δd about 0.003 and δs close to 0) and terminal substitution rates below about 0.008. Together the short fragments and the authentic but modest deamination were consistent with a genuine, degraded specimen of relatively recent, late nineteenth century age rather than deeply ancient material.

### Plastid genome-based identification

The plastome recovered from the historical specimen assembled as a circular molecule of 137,068 bp with the quadripartite organization typical of grasses, comprising a large single copy region of 81,082 bp and a small single copy region of 12,820 bp separated by two inverted repeats of 21,583 bp each, and an overall GC content of 38%. Annotation recovered the gene complement characteristic of Poaceae plastomes, including the loss of a functional *accD* and the reduction of the *ycf1* and *ycf2* genes that mark the graminid clade [50]. A near identical plastome was obtained for the reference landrace Lovászpatonai (137,066 bp). In size and structure, the historical specimen’s plastome resembled those of its Triticeae relatives, wheat (134,545 bp) and barley (136,462 bp), and was set within the range of the cereals examined, far below the pineapple [*Ananas comosus* (L.) Merr.] outgroup (159,636 bp). The annotated plastid genome is shown in Figure 2.

Comparison of the four inverted repeat junctions JLB, JSB, JSA, and JLA across nine grasses and pineapple (Figure 3) placed both rye genomes firmly within the Triticeae. As in wheat and barley, the IRb to SSC border (JSB) duplicated about 209 bp of *ndhH*, the SSC to IRa border (JSA) fell among *ndhA*, *ndhH,* and *rps15*, and the repeats extended across *trnH* and *rps19* at the JLA and JLB borders, an architecture clearly distinct from the longer repeats of pineapple.

Assembly accuracy was confirmed by amplifying and sizing the four junctions and two indel bearing regions, all matching the assembly (Figure S7). The spacer between *rbcL* and *accD* carried a 287 bp indel together with a copy of *rpl23*, whereas the functional *rpl23* lay in the inverted repeat between *rpl2* and *trnI* alongside a 285 bp indel. The copy adjacent to *rbcL* is the chloroplast pseudogene *rpl23*’, first described from wheat by Bowman et al. [79] and from rice by Shimada and Sugiura [80], where it is maintained by biased gene conversion with the functional repeat copy and predates the divergence of the cereals. We confirm its presence in rye, where to our knowledge it has not been characterized previously. Alignment against pineapple recovered the three inversions diagnostic of Poaceae, a 28 kb inversion in the *trnG* to *rps14* region, a nested 6 kb inversion in the *trnG* to *psbD* region, and a third inversion under 1 kb in *trnT* (Figure S8), showing that the historical rye plastome retained the canonical grass arrangement.

### Origin of the historical specimen

To locate the historical now identified specimen as rye within the diversity of the crop, we placed it in a maximum likelihood tree built from 16,198 genome wide SNPs across 92 samples, a genotyping by sequencing panel of 91 rye accessions spanning the cultivated subspecies rooted on *Secale strictum* and *S. vavilovii*. The specimen resolved inside *Secale cereale* subsp. *cereale* rather than with any wild or eastern lineage. Its closest relatives were cultivated accessions from Hungary (R2229) and Slovakia (R604), and it nested within a broader Central, Eastern and Northern European cluster of subsp. *cereale*. Support was consistent by SH-aLRT but weaker by ultrafast bootstrap, and the shallow backbone among cultivated accessions, expected for genotyping by sequencing data in an outbreeding crop with extensive gene flow, means the analysis placed the specimen within the European cultivated gene pool close to Hungarian and Slovak material rather than fixing a single geographic origin (Figure 4). A principal component analysis of the same SNP matrix gave a concordant placement, with the specimen falling inside the cultivated *Secale cereale* cluster (Figure 5; species-colored version, Figure S9).

### Recovery of stress associated loci from the historical specimen

Screening recovered mappable reads for 121 of the 192 reference loci, a little under two thirds of the panel. Recovery differed sharply among the three trait classes and was shaped throughout by the degraded, fragmentary nature of the template. Biotic stress resistance loci recovered best. Reads mapped to 28 of 34 references at a median maximum depth near 10x, and 17 loci carried at least one substitution relative to the reference. Recovery reached the nematode resistance Cre3 homologues, the Sr50 (RGA1) stem rust locus, NBS-LRR resistance gene analogues, the powdery mildew ARM1 and PUB15 genes, and the acetolactate synthase herbicide target, so the principal resistance gene families of rye were represented in the specimen. Abiotic stress references mapped most often, 39 of 42, but shallowly. Median depth was about 6x and 16 loci sat at or below 5x, so presence was clear while depth was marginal. Consensuses were almost entirely identical to the reference, consistent with conspecific material and with limited variant information at low depth. The aluminum tolerance system behaved differently; only 54 of 116 references recovered a consensus. The many short-tagged sites and random amplified marker sequences of the Alt3 and Alt4 loci frequently returned nothing, whereas the ALMT1 cluster and the ScAACT1 (ScMATE) genes recovered in part. The low rate here reflected reference composition, since marker amplicons and long complete coding sequences are the least favorable targets for short-degraded reads. The signature of fragmentation ran through every class; of the 121 recovered loci, 77 came back not as a single continuous consensus but as two or more separate blocks. Long references and full-length coding sequences were routinely reconstructed only in patches, with internal gaps, and the least complete recoveries fell away to no consensus at all, which accounts for most of the 71 references that failed. Maximum depth was modest across the panel and fell towards the lower categories, so many calls were resting on thin coverage. Read against these limits, stress-associated loci of rye remained present and recoverable in the historical specimen. It does not, however, and given conspecific material cannot, demonstrate that any trait was functionally active. Per locus results are given in Supplementary Material 3 and summarized in Figure 6.

Beyond the stress panel, we examined the three heading date QTLs of rye. At Hd2R, the ScPpd1-associated locus on chromosome 2R, the site was covered deeply, and the specimen carried the Weining early heading allele without ambiguity, with a cytosine read on 2635 of 2742 reads and an even split across strands. The other two QTLs recovered only five to six reads each. At both Hd5R and Hd6R the majority base differed from the Weining earliness allele, tentatively pointing to the alternative state, but at Hd5R the difference is of the C to T class produced by post mortem deamination and at Hd6R the supporting reads fell on a single strand, so neither genotype could be fixed at this depth (Table 1). One of the three earliness determinants was therefore present in the specimen, while the remaining two stayed beyond reliable recovery.

**Table 1.**
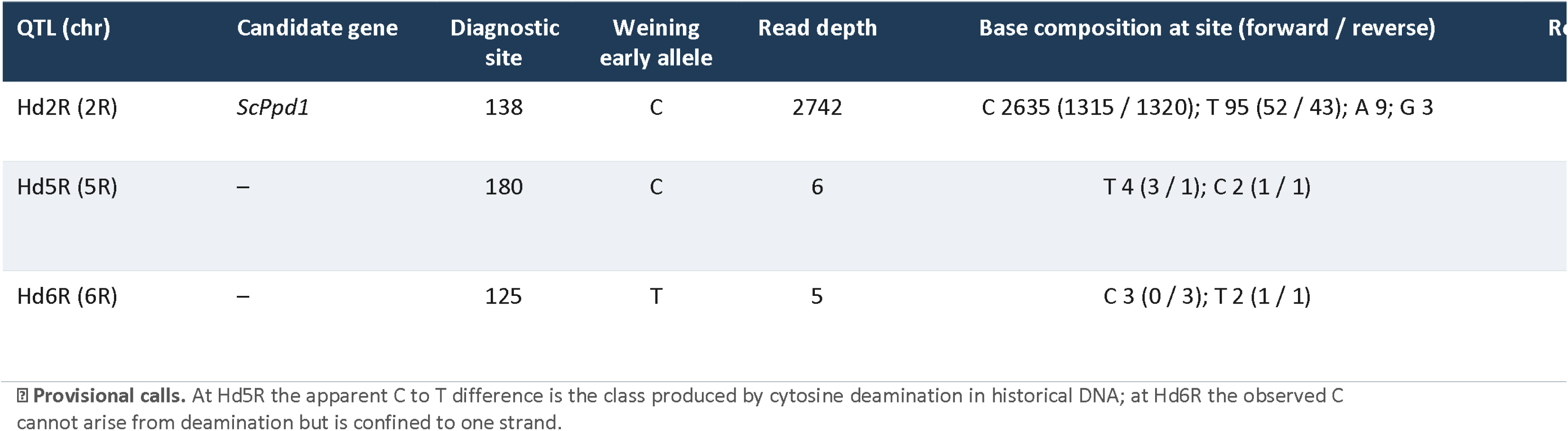
Genotyping of three rye heading date QTLs in the historical specimen against the Weining reference.

### Divergence of modern rye from the historical landrace at selective sweep loci

We next asked how far modern rye has diverged from the historical landrace at loci implicated in domestication and improvement, using the 36 candidate selective sweep regions that Li et al. [58] detected in the Weining genome (flagged by three sweep statistics, DRI, XP, and FST; comprising 13 DRI, 18 XP, and 5 FST regions) and totaling about 219 kb (Supplementary Tables S2 to S4). Within these regions the old rye differed from the Weining reference at 3231 sites, corresponding to 1.48% of the sequence examined, while the three whole genome resequenced cultivars differed from the old rye consensus at 3636 sites for R2446 (1.66%), 3052 for R925 (1.39%), and 1187 for PUMA (0.54%). PUMA was thus the modern cultivar most similar to the historical rye across these sweep loci, and R2446 the most divergent, differing from it about three times as often as PUMA (Figure 7). Divergence was distributed unevenly among the regions, with a small number of loci accounting for most of the differences while many carried only a handful, so the signal reflected a few strongly differentiated domestication loci rather than a uniform shift. Resolving the divergence by the signal that identified each locus sharpened this picture. PUMA remained the least diverged line in every class (0.32% in DRI, 0.90% in XP, and 0.26% in FST regions), whereas the other lines reordered by class. In the DRI regions the Weining reference was itself the most diverged from the old rye at 1.55%, ahead of R2446 (1.10%) and R925 (0.72%), as expected for loci defined by a reduction of diversity on the Weining genome, while in the XP regions this ordering reversed, with R2446 and R925 the most diverged at 2.43% and 2.36% and Weining intermediate at 1.55%, and the five FST regions followed the same ordering at a lower level. The historical rye therefore diverged most sharply from present day cultivars at the XP sweep loci, while retaining closest similarity to PUMA throughout.

## Discussion

The main objective of this study was to identify the recovered cereal specimen and probe its origin by genome skimming, rather than to reconstruct a complete nuclear genome. Historical plant material is characteristically fragmented and depleted, and its endogenous DNA is often too degraded for reliable whole genome assembly [6]. Genome skimming circumvents this by sequencing total DNA at low coverage and recovering the naturally high copy fraction, the plastid and ribosomal genomes together with a broad, shallow sample of nuclear reads, from exactly the low-quality input that defeats targeted approaches [9,12]. Such data are sufficient not only for confident taxonomic assignment and phylogenetic placement [10,11], but also for scanning defined nuclear regions. Let us interrogate the ancestry of the historical rye and its relationship to modern varieties beyond a single barcode.

### The historical rye at the threshold of modern breeding

The assembled plastome placed the historical sample securely within *Secale*, and its nuclear genome wide profile within cultivated *Secale cereale* subsp. *cereale*, closest to a Hungarian bred rye and a Slovak upland landrace and remote from any wild or eastern lineage. The placement of the sample gains full meaning when read against the agricultural history of the region. R604, one of the identified closest accessions to our sample, is a tall winter landrace collected at Močiar in the southern foothills of the Slovak Ore Mountains at 760 m. According to its Gatersleben evaluation recording, the plants are 1.6 to 1.9 m tall, with sparse rust and smut infections occurring on them, typical of those traits of a hardy upland rye [81]. While R2229, the other accession closest to the historical samples, is not a landrace but a Hungarian variety registered at the same collection as Hungarian Giant [81]. Our sample was recovered in the same country with typical tall features; thus, genomic features, geography, and stature confirmed that it belonged the Carpathian Basin gene pool and its Slovak margins point to a specific episode of rye production. Through the closing decades of the nineteenth century, the rye of Central Europe was reshaped by the first systematic selection from landraces, and by about 1880 intensified breeding had yielded taller, more productive, and more lodging resistant populations that spread quickly across the region [24]. Hungarian farmers met this wave of new varieties as reported in a county newspaper of 1891, which pressed its readers to sow the newest improved rye:

In recent years the rye crop has been enriched with several excellent new varieties, yet all are surpassed by the two newest, the Schlanstedt giant rye and the sack filling Mammoth rye. Both possess every quality a good rye should have; the former, in fair weather, yields as much as fifteen hundred kilograms to the cadastral yoke, while the latter earned its name through sheer abundance, and Ödön Mauthner hastened to procure both for farmers who value progress. [82]

The described stature ‘Mammoth’ is quite telling regarding our samples, but the Schlanstedt giant rye of that notice is not unknown for rye breeders either. Schlanstedt was the estate of Wilhelm Rimpau (1842–1903), whose improved Schlanstedt variety supplied the pollen parent of the first fertile amphidiploid of wheat and rye in 1888, and whose account appeared, as it happens, in that same year of 1891 [83]. His Schlanstedt population, with von Lochow’s Petkus rye, stood at the head of the improved Central European ryes, and the Hungarian source shows one of them was offered to farmers. It is plausible that such a variety was in cultivation in the region and the straws were used to fill in the settee of the *canapé* at a time. On the other hand, the tall Hungarian and Slovak ryes among which the historical samples fell, preserved in the IPK collection, can be regarded as the living and lexical residue of the rye improvement momentum.

### What the recovered loci reveal about the old rye

The loci recovered from the historical sample sketch a trait repertoire spanning three axes, defense against pests and pathogens, protection against abiotic stress, and tolerance of mineral toxicity. Among the resistance references we recovered homologues of the Cre3 receptor-like kinase family linked to nematode resistance [84], the Sr50 stem rust locus [85], the ARM1 and PUB15 U-box armadillo E3 ligase genes that confer powdery mildew resistance in the Triticeae [86], and several NBS-LRR resistance gene analogues. We also recovered acetolactate synthase, an essential housekeeping enzyme of branched chain amino acid biosynthesis that only much later became the target of ALS inhibiting herbicides [87]. The recovery here reflects the conserved gene body alone, since the resistance conferring mutations reported were selected under herbicides in the late twentieth century and postdate this specimen by well over a century. With that qualification, the specimen held the principal disease and pest resistance gene families known from modern rye. On the abiotic side we recovered the alkylresorcinol synthase gene, which builds the phenolic cuticular wax that shields rye against desiccation, ultraviolet light and pathogen ingress [88], along with several rye stress responsive transcripts. Most striking was the aluminum tolerance system. We recovered parts of the ALMT1 malate transporter cluster and its associated Alt4 markers [89–91], the module that lets rye secrete malate and neutralize toxic Al^3+^ at the root tip. Because rye is the most aluminum tolerant of the small grain cereals, the presence of this system fits an old landrace adapted to acidic, aluminum rich soils.

These inferences need cautions interpretation since recovering a homologue shows only that the locus was present and recoverable in this genome. It does not show that the gene was transcribed, that a functional allele was carried, or that the plant expressed the phenotype, and post mortem damage with low depth further blunts any functional reading [60]. The claim that we can make here remains narrower yet worthwhile; the specimen preserved the structural machinery of the resistance, stress and aluminum tolerance pathways of rye, and at those loci where the recovered variants align with known tolerant haplotypes, they point, tentatively, towards real adaptive potential.

A comparable reading applies to flowering time. Rye landraces were historically later and more photoperiod sensitive than the elite varieties that displaced them, and the shift towards early, day length independent heading is among the clearest signatures of rye improvement [92–94]. Against this background the recovered specimen is quietly informative. At Hd2R, the locus over the photoperiod gene ScPpd1, it carried the same early heading allele that is fixed in the Weining rye [58], so part of the earliness architecture of the crop was already present in our historical material. The two remaining heading date loci, Hd5R and Hd6R, yielded too few reads to genotype with confidence, and where our specimen seemed to depart from the modern allele the change fell in precisely the configurations most exposed to post mortem damage. Therefore, we can only make a partial conclusion; one of the three earliness determinants was demonstrably in place, while the others remain open. The occurrence of the early allele of ScPpd1 in this historical rye accession is particularly intriguing when compared with the history of the early, insensitive allele in the orthologous Ppd-D1 gene in wheat, which has the largest effect among all the photoperiod response genes [95,96]. Historical pedigree and haplotype analyses indicate that the photoperiod-insensitive allele originated in East Asian, most likely Japanese, germplasm and was introduced into European wheat breeding only during the early twentieth century through the incorporation of Japanese breeding material [97–99]. Thus, whereas the photoperiod-insensitive allele in wheat represents a relatively recent introduction into European breeding, the presence of an early ScPpd1 allele in a historical rye accession suggests that variation promoting early flowering may have been present in European rye populations considerably earlier.

### Divergence at selective sweep loci

Selective sweeps typically reduce diversity in the corresponding region, creating long stretches of the genome with unusually few SNPs [100]. By comparing the rate of polymorphism between the historical rye and modern varieties, we obtained a snapshot of the putative sweeping process, observing the increasing homogeneity of these regions across time. The 36 regions examined here were identified by Li et al. [58] as bearing signatures of domestication and improvement in the Weining genome, and our finding that divergence from the historical rye concentrates in a small number of loci, while many regions carry only a handful of differences, is consistent with localized rather than genome wide change.

The most interpretable pattern is the reordering of lines between sweep classes. In the DRI regions Weining itself was the most diverged from the historical rye, which is expected because those loci were delimited by a reduction of diversity measured on the Weining genome and therefore carry an ascertainment bias towards Weining specific variation. The XP regions behaved differently, with R2446 and R925 diverging most strongly and Weining intermediate. Because cross population statistics identify loci differentiated between germplasm pools rather than depleted within a single genome, the XP loci appear to be where improvement germplasm has moved furthest from the landrace state, and where the historical specimen retained a configuration closer to the pre breeding condition.

PUMA was the least diverged line in every class. It may share a comparatively recent ancestry with the germplasm from which the historical rye derives, or its swept regions may have been fixed for alleles already present in the landrace, in which case the low divergence records the retention of older variation through breeding. Distinguishing these possibilities requires allele frequency data from several historical individuals, and this is the main limitation here, since pairwise divergence from a single genome measures difference rather than diversity and cannot separate selection from drift.

Historical landrace collections have been shown to contain a substantial amount of novel genetic diversity that has since been lost through previous selection sweeps, in wheat [101,102] and in rye and barley [16,18], and plastid data from historical rye point the same way [24]. Because rye was domesticated from weedy progenitors and improved intensively only within the last two centuries [103,104], swept loci are precisely where recent selection should be visible, and historical specimens offer a direct test of when that variation was lost [6].

## Conclusions

Our study demonstrates that a single degraded historical specimen, analyzed through damage-aware sequencing and interpreted alongside a modern reference genome, genebank passport data, and contemporary documentary evidence, can be resolved simultaneously to a gene pool, a constellation of agronomic traits, and a specific moment in agricultural history. The cereal recovered from late nineteenth-century furniture is best understood as a tall Central European rye, occupying the transitional space between traditional landraces and the emerging improved varieties, and likely bearing the imprint of the Schlanstedt and Mammoth ryes that were rapidly spreading across the region at the time. The genetic evidence is remarkably consistent across every line of inquiry. The specimen falls unequivocally within cultivated *Secale cereale* subsp. *cereale*, showing no affinity with wild or eastern lineages. Across domestication sweep regions and loci controlling heading date, it already possessed elements of the modern genetic architecture for earliness, including the Weining allele at *Hd2R* associated with the photoperiod response gene *ScPpd1*. Yet, as expected in an obligately outcrossing crop, the shallow SNP structure among cultivated accessions does not permit assignment to a single named variety. Instead, the historical specimen is most appropriately placed within the broader Central European breeding population from which improved cultivars were being assembled. We believe, this study demonstrates that historical crop remains, when examined through an integrated genomic and historical framework, preserve not only genomes but also the material traces of human agricultural practice. In our case, these traces illuminate the gradual transition from locally adapted landraces to scientifically bred cultivars, revealing how the biological and cultural histories of agriculture became inseparably entwined.

## Acknowledgements

The authors gratefully acknowledge Antal Balázs, a conservator from Rákóczifalva, Hungary, who discovered the historical rye sample while restoring the late nineteenth century antique settee, and kindly provided it for this study. We also thank Elina Laiho from the Historical DNA laboratory of LUOMUS for assistance. We thank the CSC – IT Center for Science, Finland, providing free access to high-performance computing resources essential for conducting this research in support of academic study and education. We also thank the Helsinki University Library for supporting open access publication. The authors sincerely thank Dr. Christina Hartmann (Department of Genebank, Research Group *Resources Genetics and Reproduction of the Rye Germplasm*, Leibniz Institute of Plant Genetics and Crop Plant Research (IPK), Gatersleben, Germany) for generously sharing her expertise, historical information, and photographs, which were invaluable in interpreting the historical context of this study. AC thank the support of the János Bolyai Research Scholarships of the Hungarian Academy of Sciences (BO/00416/23/4). We thank Jennifer Rowland for editing the early version of the manuscript.

## Data Availability

Short reads are available under BioProject PRJNA1510302, and the chloroplast genome assemblies are available in NCBI under PZ895511-PZ895512.

## Contributions

P.P., T.N., O.V., I.K. and A.Cs. conceived the study. P.P. designed the methodology and led the bioinformatic analysis and validation, with further contributions from S.J., S.W. and R.D. Formal analysis was carried out by P.P., S.J., Z.K. and R.D., and the investigation and data generation were performed by P.P., S.J., Z.K., L.C., R.D., T.N. and A.Cs. Resources were provided by P.P., T.N., I.K. and A.C., while Z.K. and S.W. curated the data. P.P. wrote the original draft, and S.W., R.D., I.K. and A.Cs. contributed to reviewing and editing the manuscript. Visualisation was prepared by P.P., S.J., L.C. and R.D. P.P. and O.V. supervised the work, P.P. administered the project, and P.P. and T.N. acquired the funding. All authors read and approved the final manuscript. † Tamás Németh passed away during the preparation of this manuscript. The authors dedicate this work to his memory.

## Declaration on the use of generative AI and AI-assisted technologies

During the preparation of this work, we used Claude (Anthropic) Opus 4.8 with *High* effort setting to reformat the in-text citations and reference list into *Scientific Reports* style and to check spelling and grammar. After using this tool, we reviewed and edited the output and take full responsibility for the content of the publication.

## Supplementary Information

***Supplementary Material 1*** – Tables and figures.

***Supplementary Material 2*** – Nuclear SNP matrix use for phylogenetic analysis.

***Supplementary Material 3*** – Trait assessment by read data-mining.

---

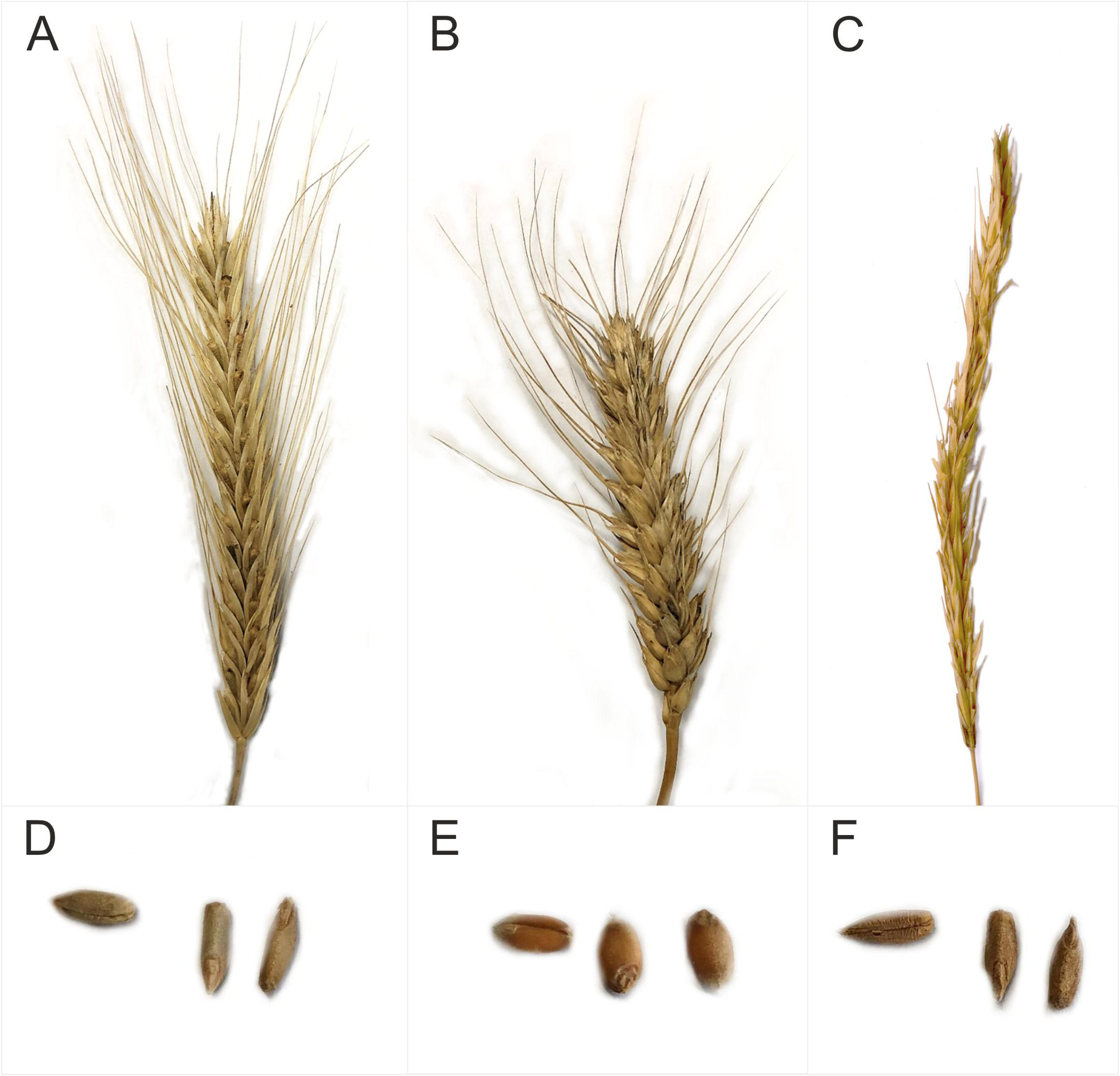

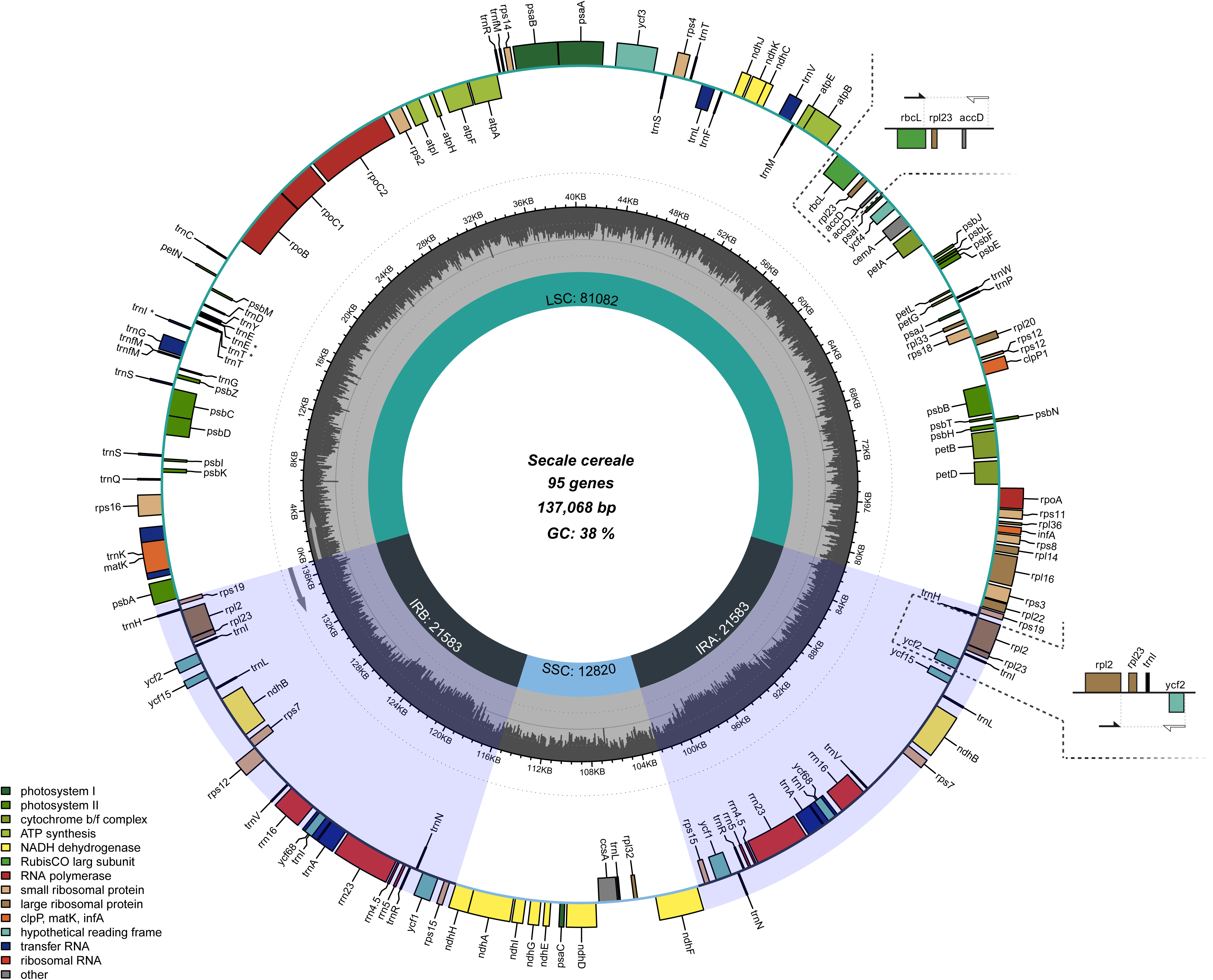

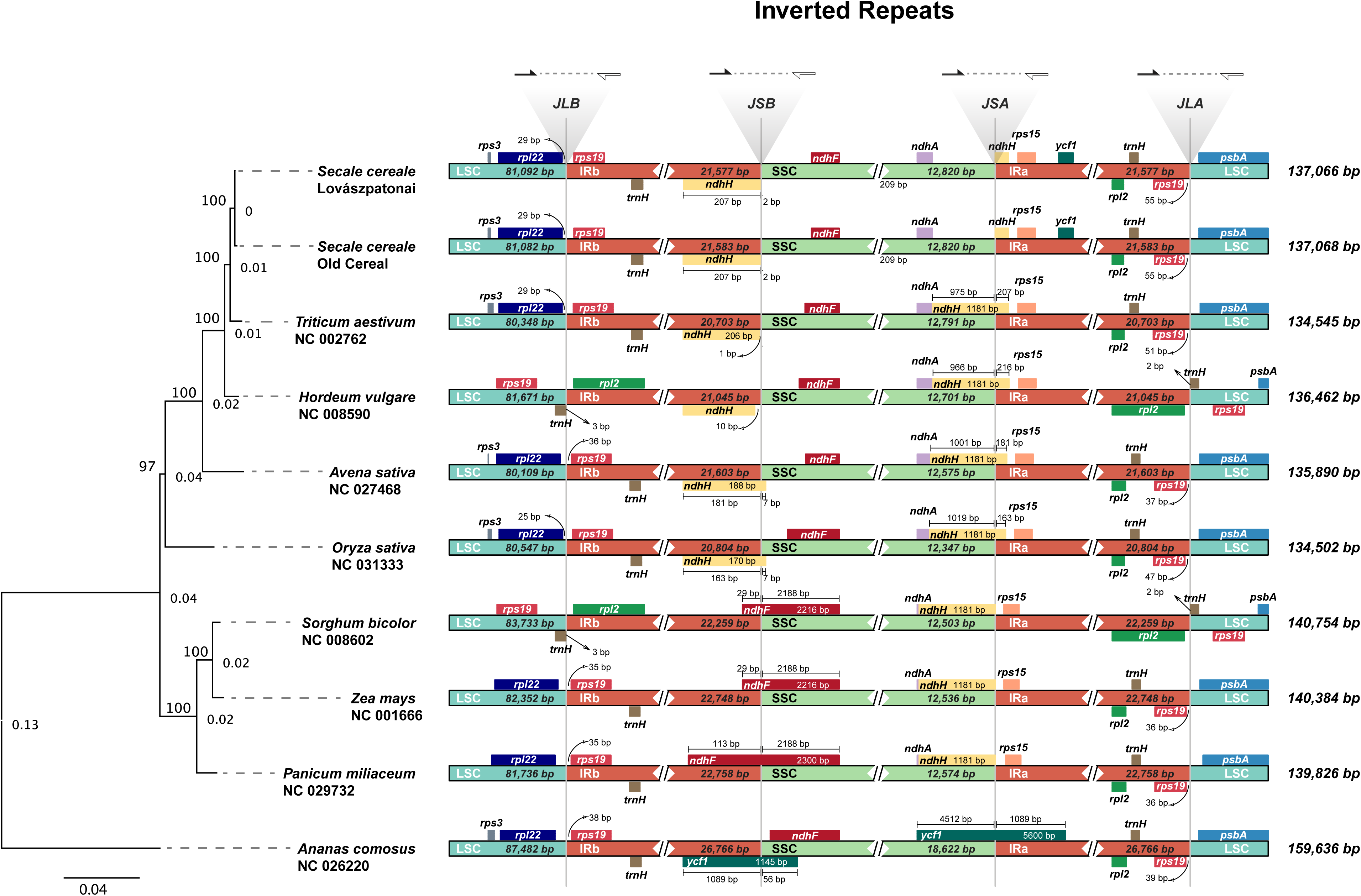

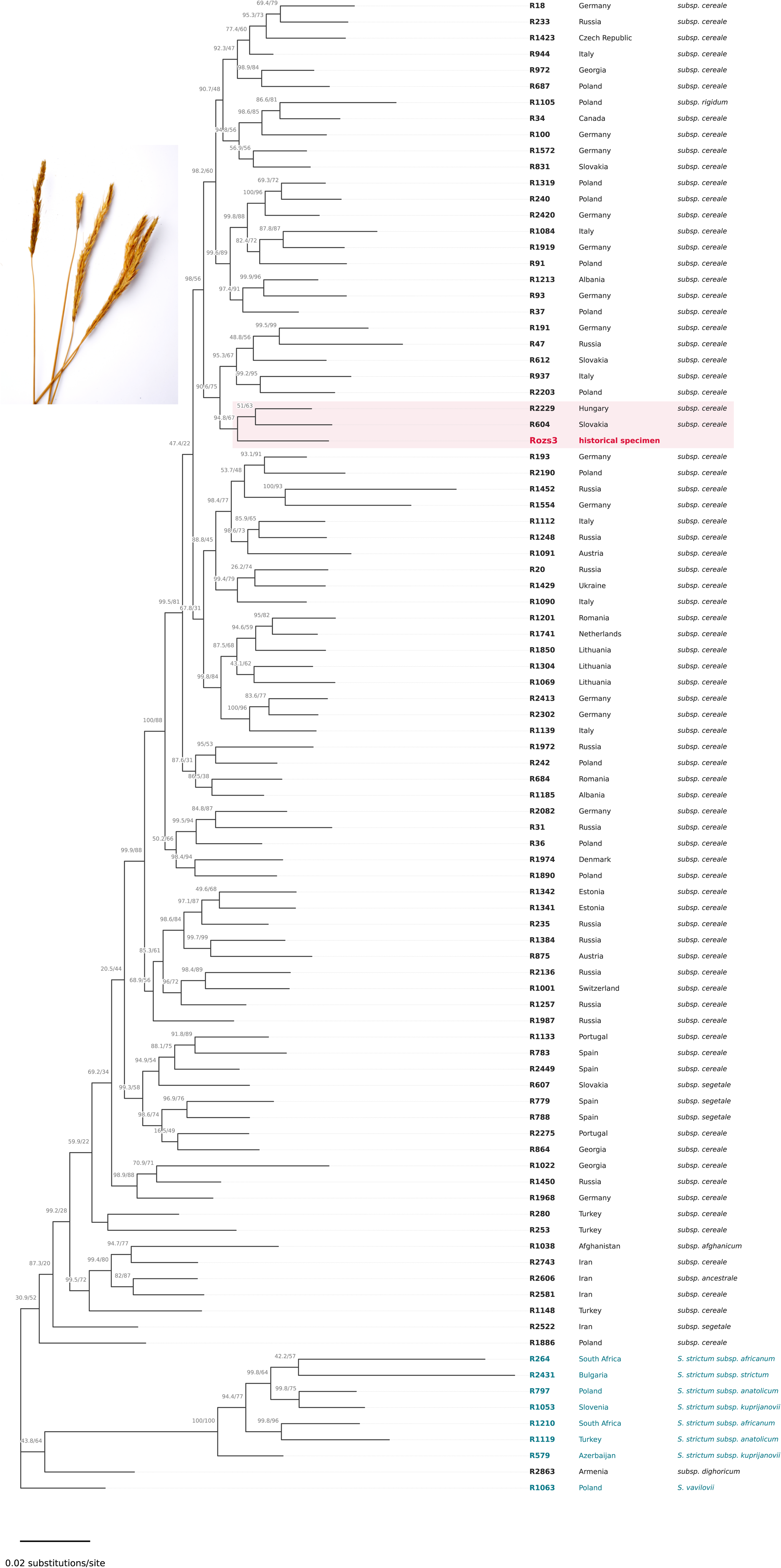

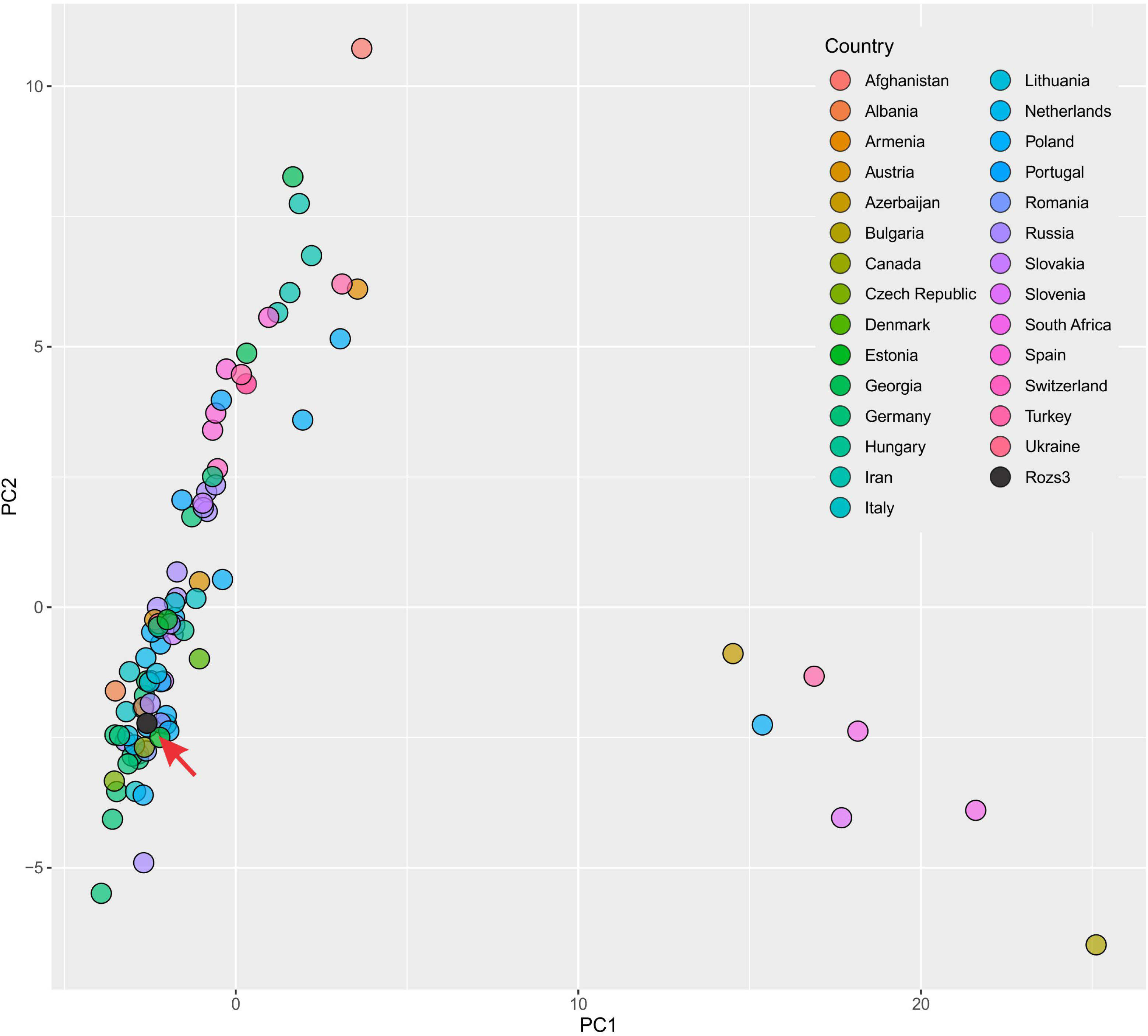

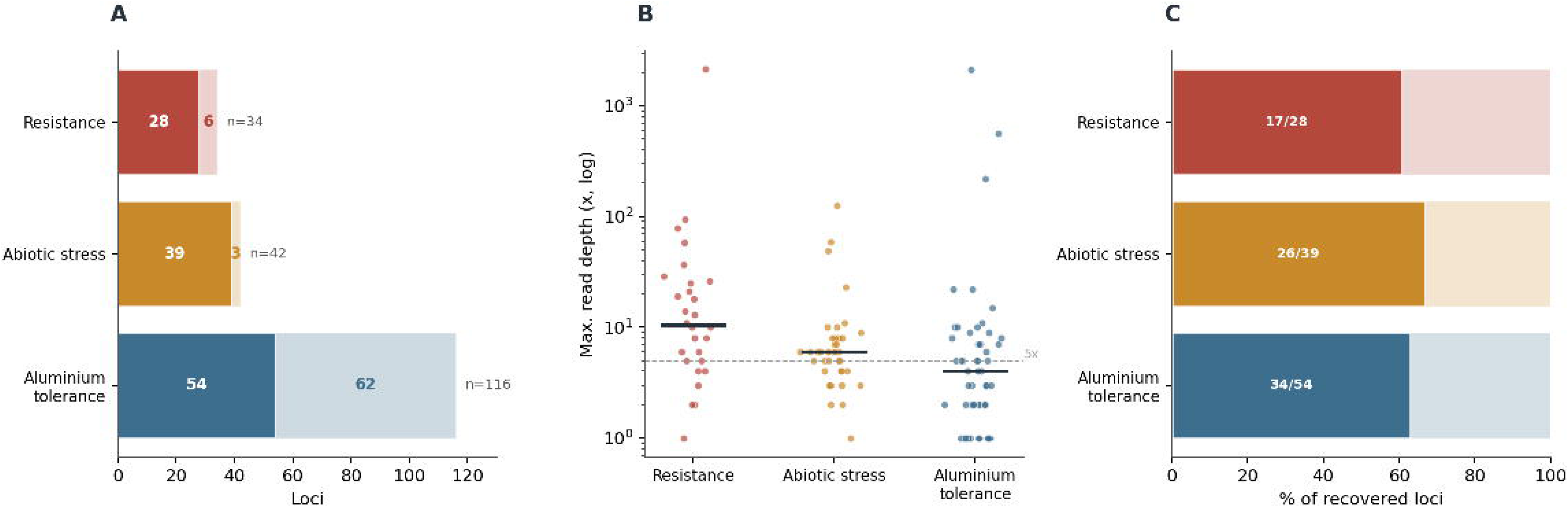

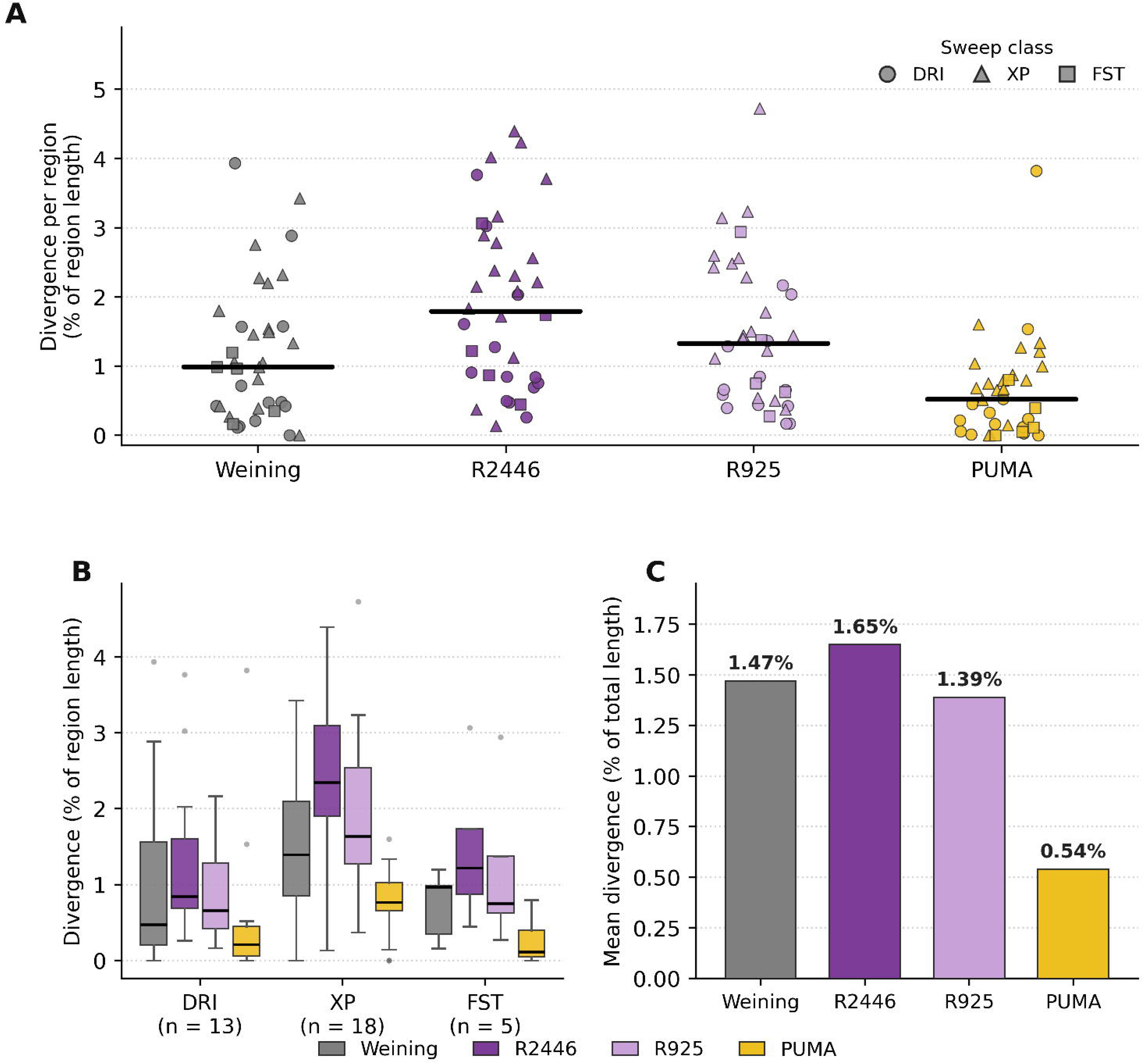

