## Supplementary Material 1 for "Catcher in the rye: museomics reveals the identity of cereal remains recovered from a late nineteenth century antique furniture"

*Supplementary figures and tables*

**Supplementary Figures**

**(A)**

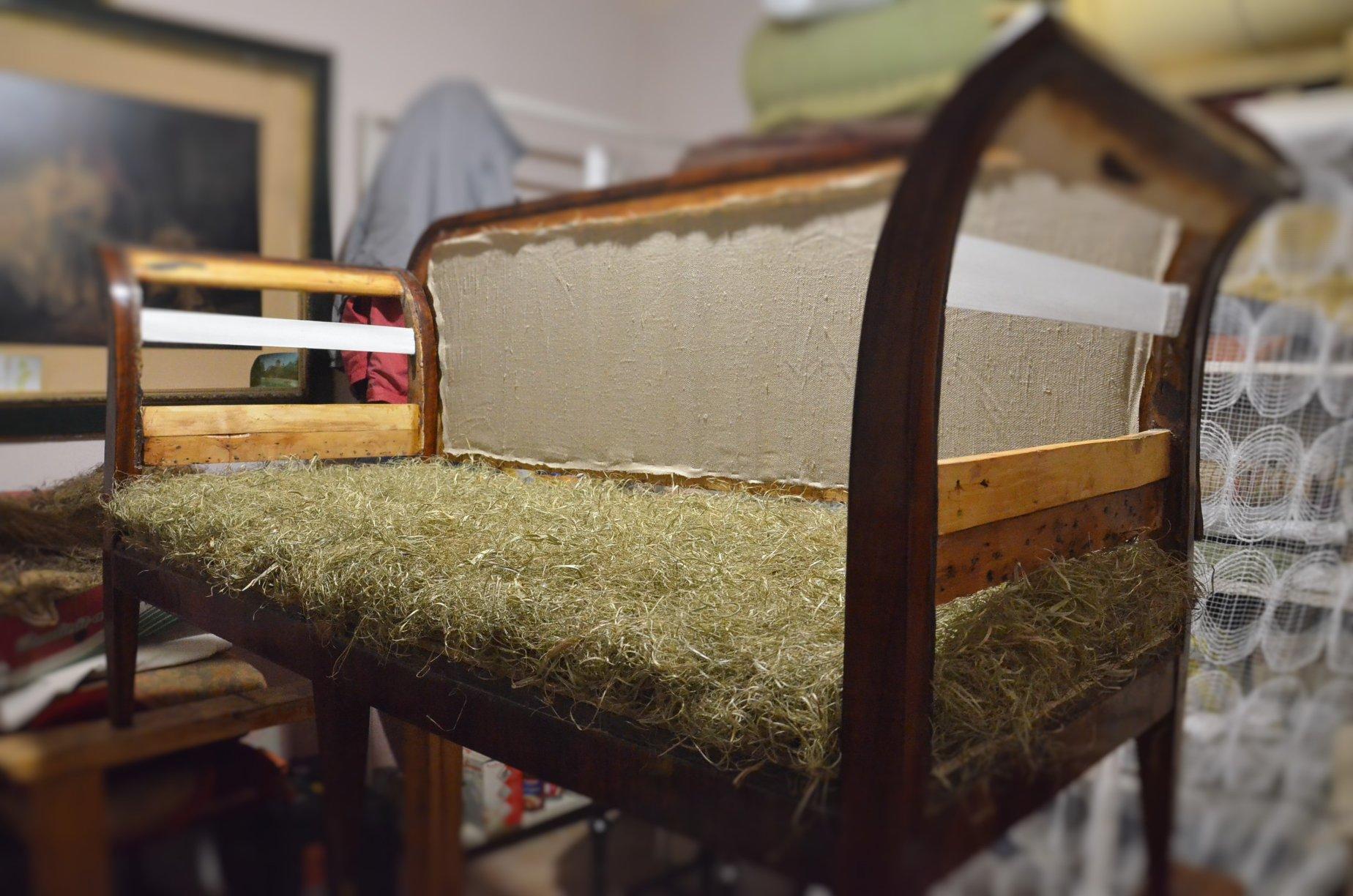

**(B)**

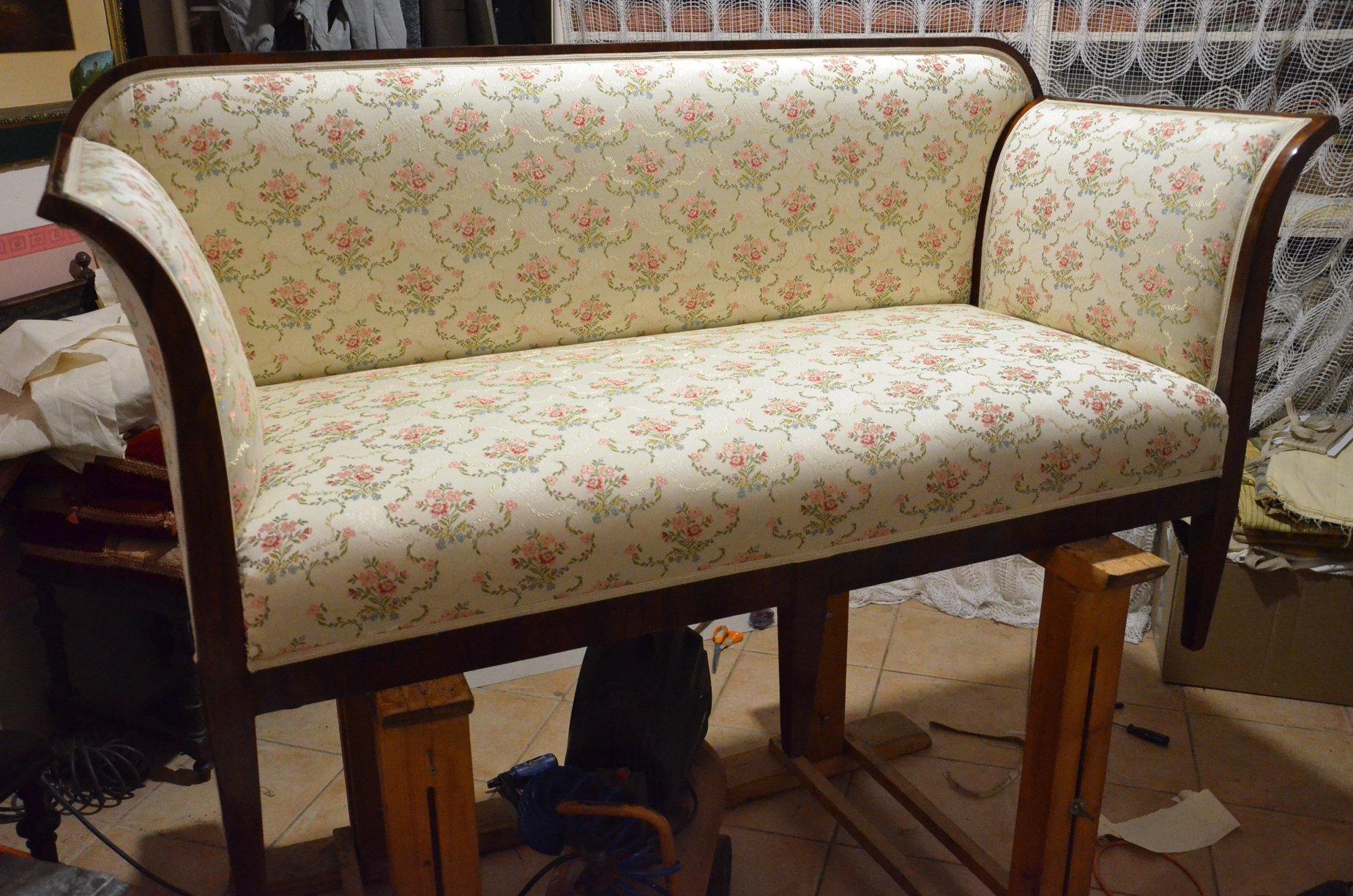

**(C)**

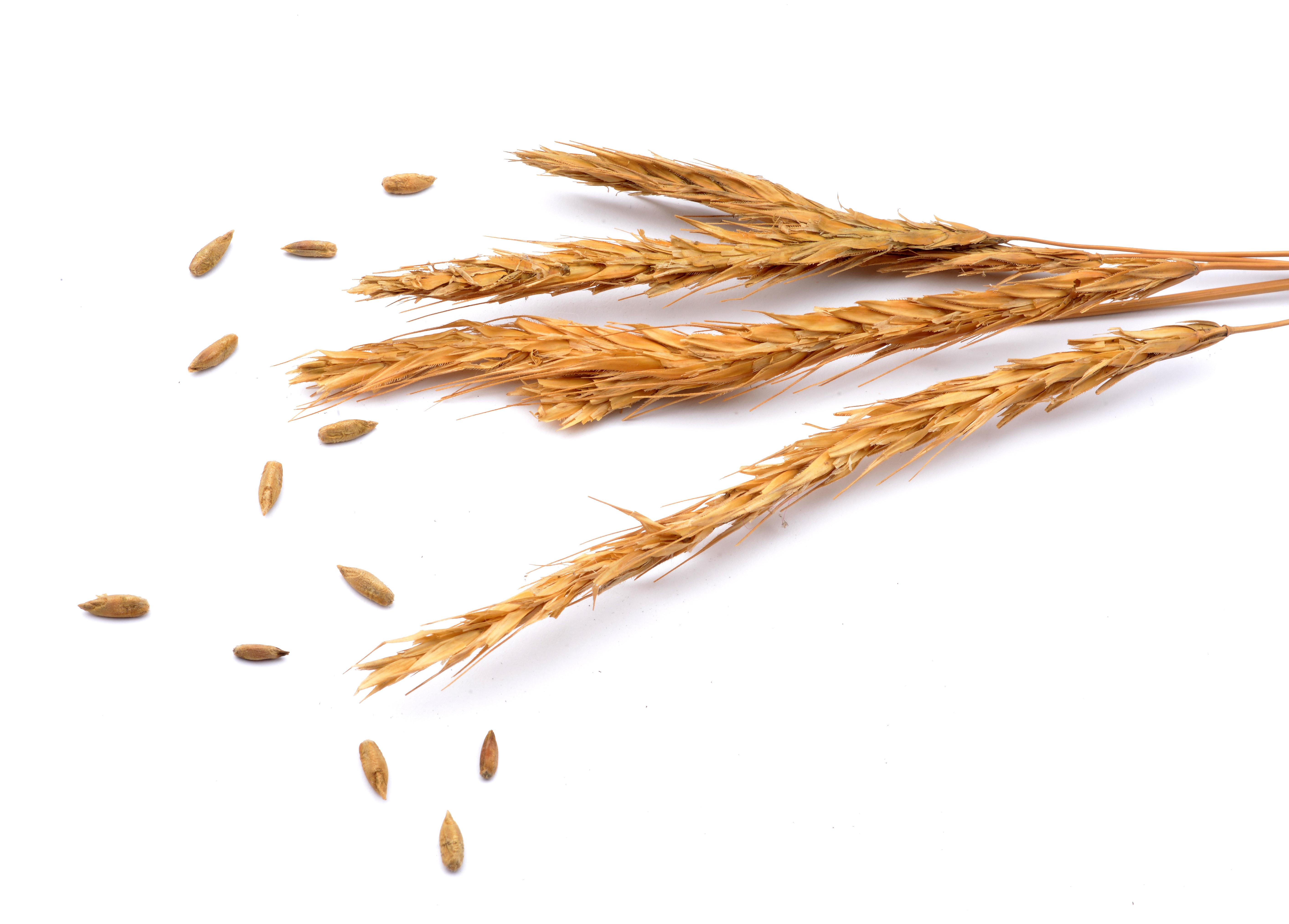

**Figure S1. The historical specimen and its recovery context.** The cereal was recovered as upholstery padding from an antique settee, assessed as a canapé in the Habsburg Biedermeier manner, more precisely a later Biedermeier revival of the Historicist or Gründerzeit period dating from about 1870 to 1900. (A) The piece during conservation, with the original rye seat padding exposed beneath the hessian and linen cover. (B) The same canapé after restoration and reupholstery. (C) Rye ears and loose caryopses recovered from the original upholstery. At the time of sampling the original layers were undisturbed, a sealed context that minimises modern contamination and supports the historical integrity of the material.

**Post-mortem damage assessment (mapDamage2.0)**

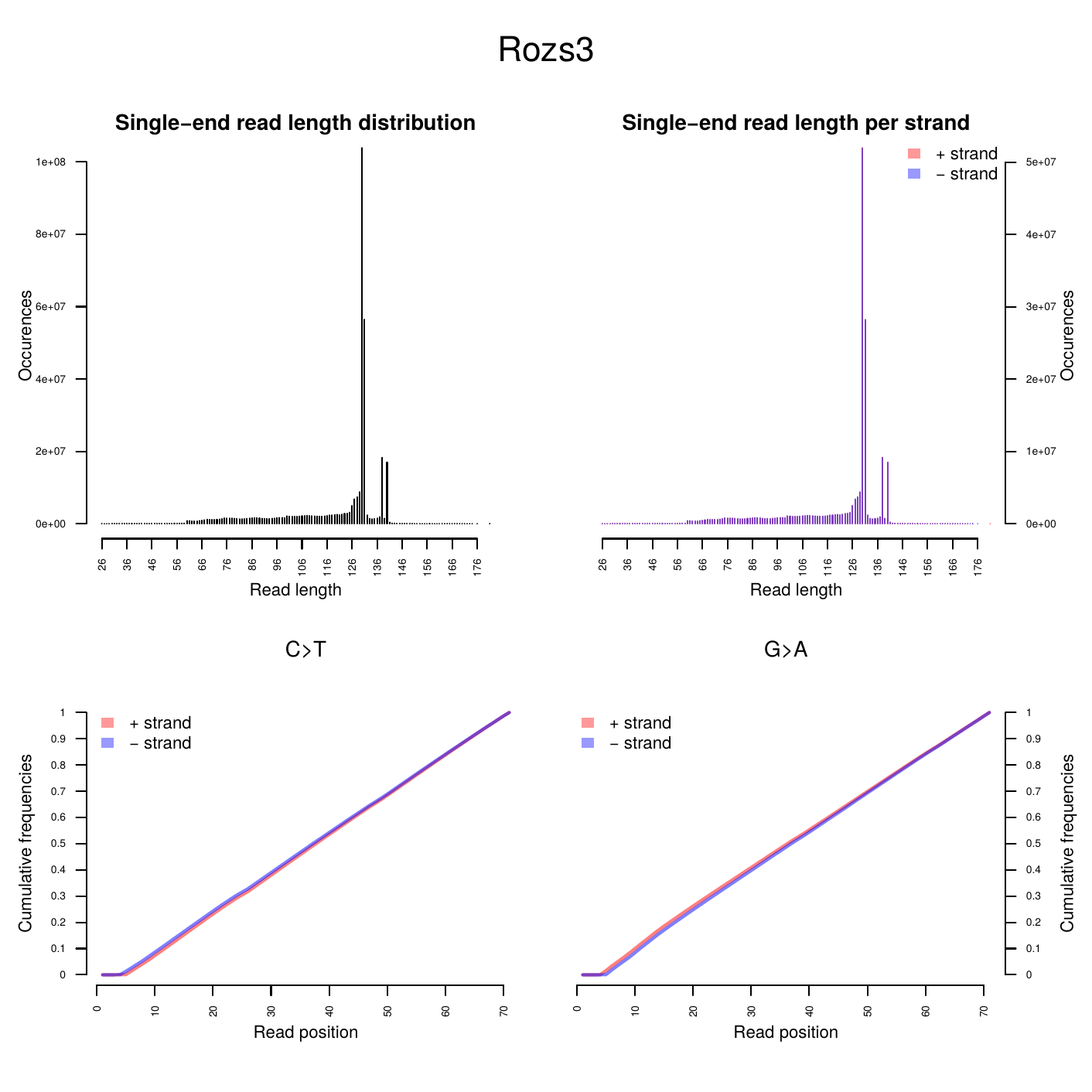

**Figure S2. Fragment length distribution of the historical specimen (Rozs3) sequencing reads, computed with mapDamage2.0.** The left panel shows the single end read length distribution and the right panel the distribution separated by strand (plus and minus). The concentration of reads at the shortest lengths, declining toward longer molecules, is characteristic of degraded historical DNA.

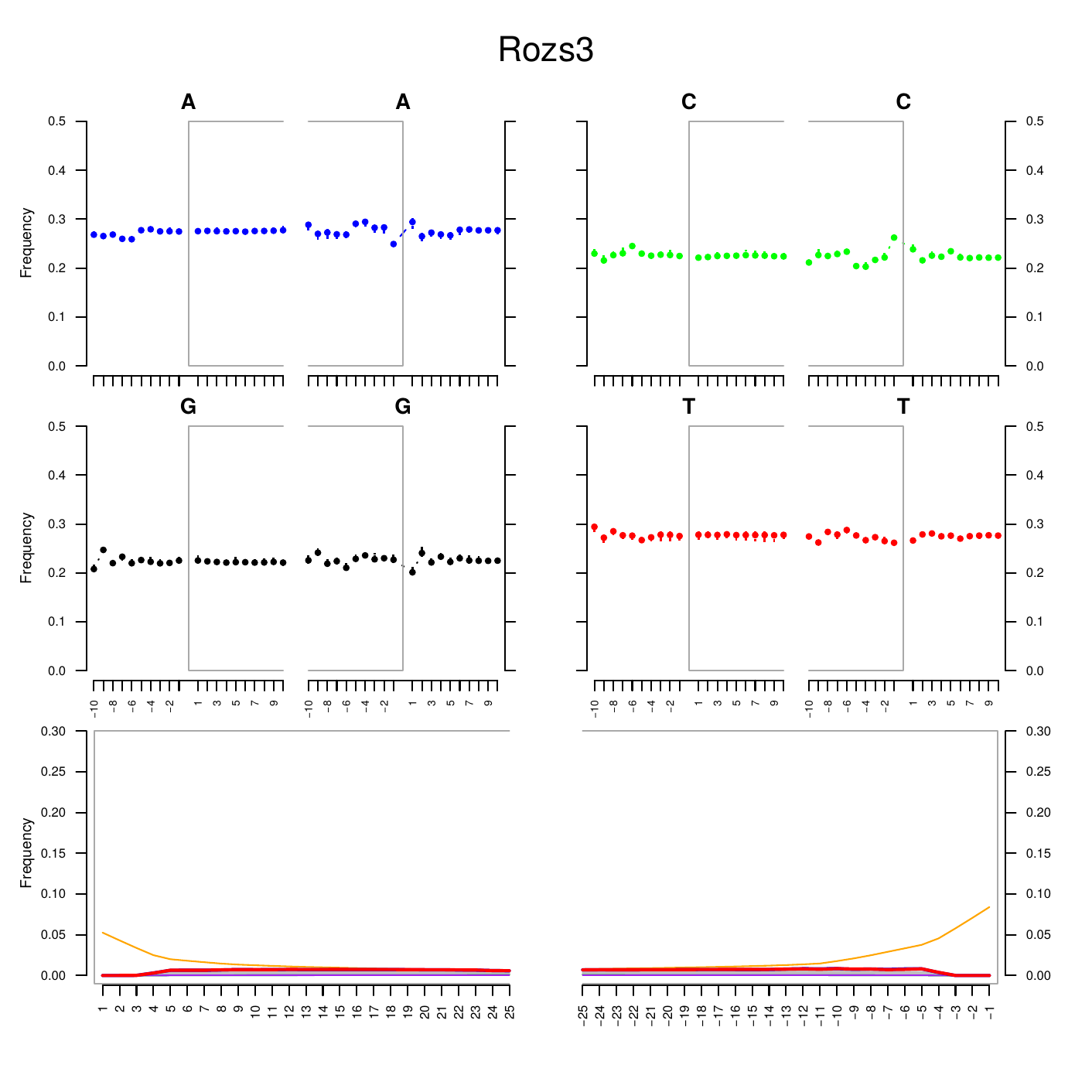

**Figure S3. Nucleotide misincorporation and base composition around read termini for the historical specimen (Rozs3), from mapDamage2.0.** Upper panels show base frequencies in the reference regions flanking the reads. Lower panels show the per position substitution frequencies, with C to T substitutions (red) elevated toward the 5′ end and complementary G to A substitutions (blue) elevated toward the 3′ end, the diagnostic signature of post mortem cytosine deamination. Grey lines show all other substitution types.

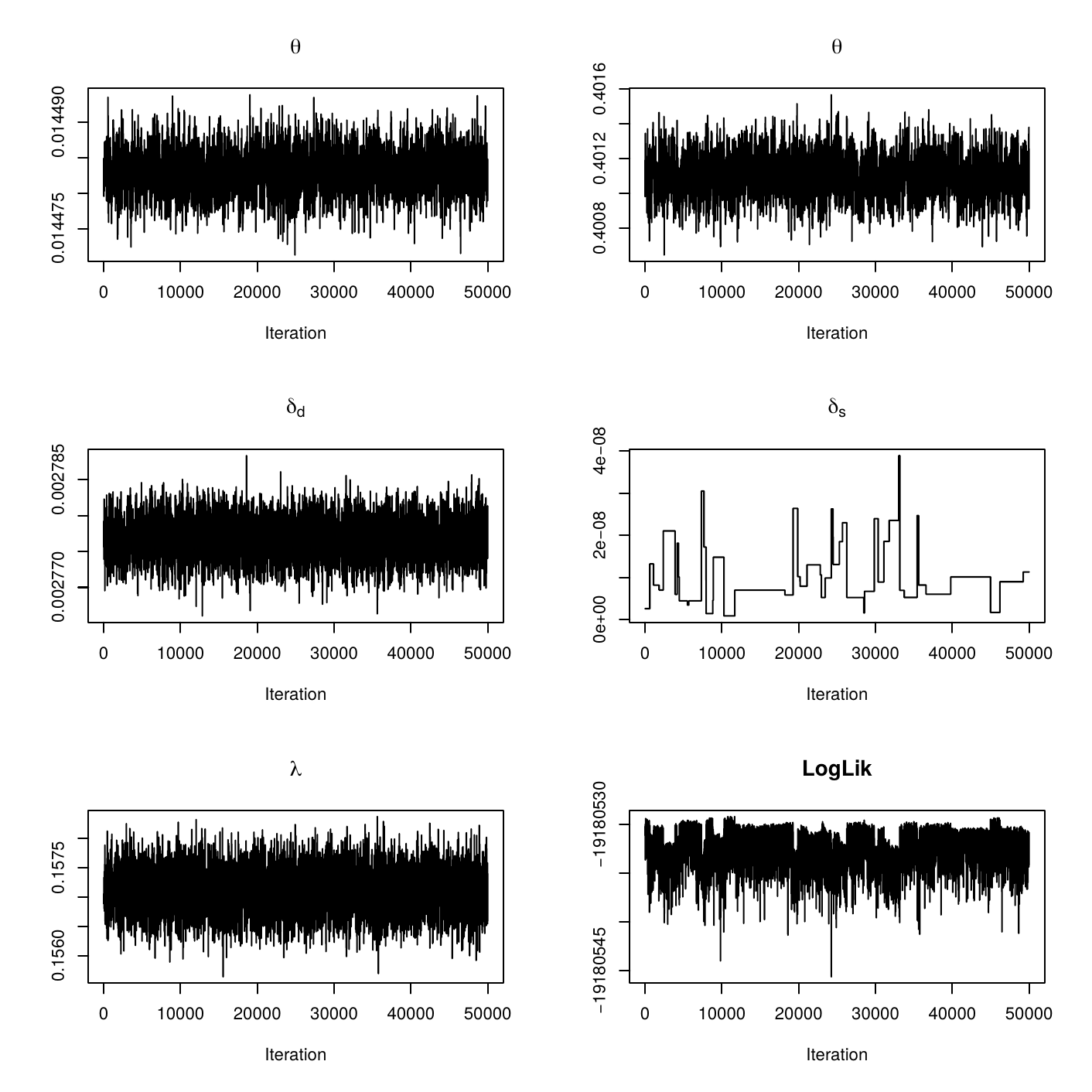

**Figure S4. Trace plots of the mapDamage2.0 Bayesian damage model parameters across 50,000 MCMC iterations for the historical specimen (Rozs3).** Stable, well mixed chains for the overhang length parameter λ, the double and single stranded cytosine deamination rates δd and δs, the substitution parameters θ and ρ, and the log likelihood indicate convergence of the sampler.

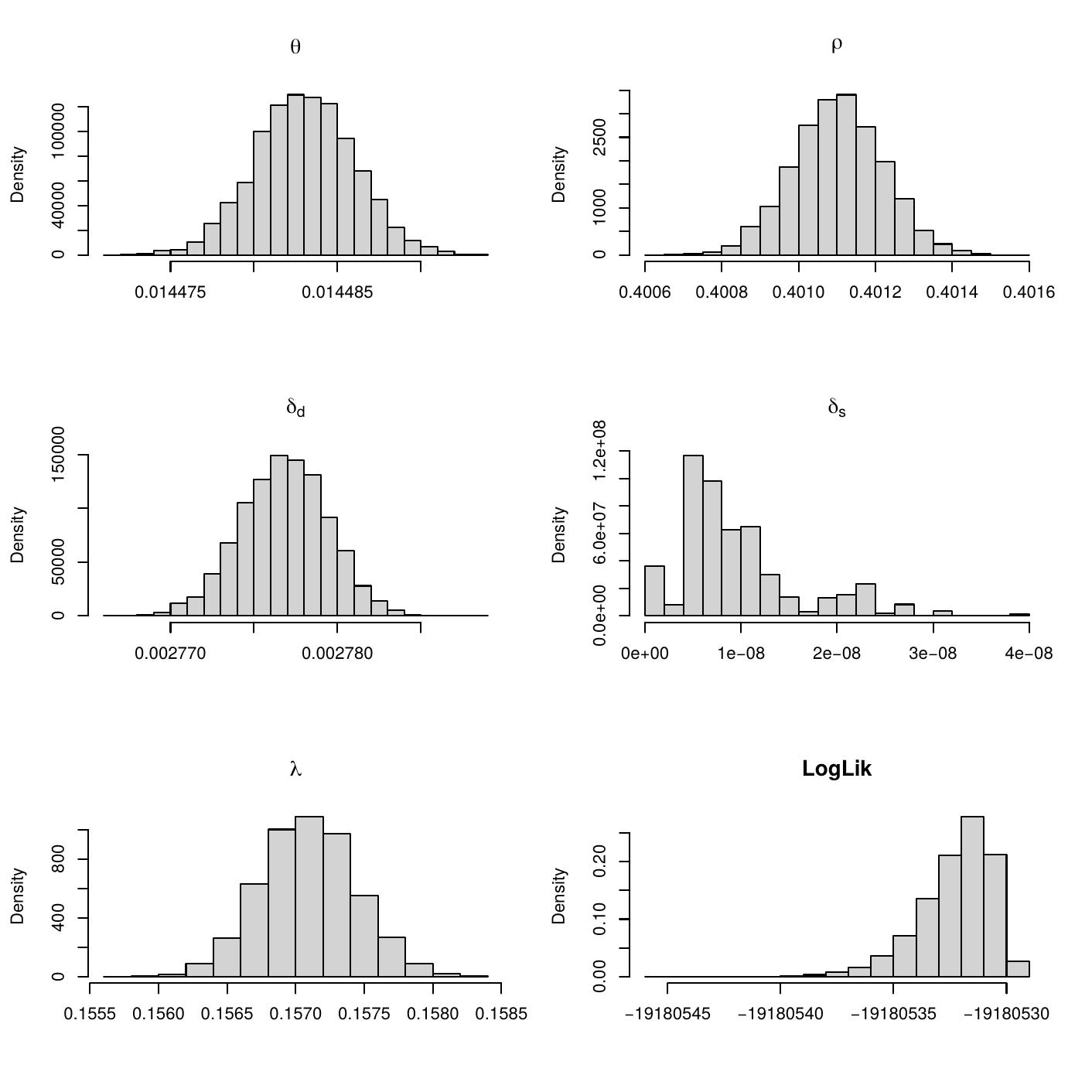

**Figure S5. Posterior distributions of the mapDamage2.0 damage model parameters for the historical specimen (Rozs3), summarizing the same MCMC run as Figure S4.** The narrow posteriors, with λ near 0.16 and low deamination rates (δd about 0.003 and δs close to zero), indicate limited post mortem damage and precise parameter estimates.

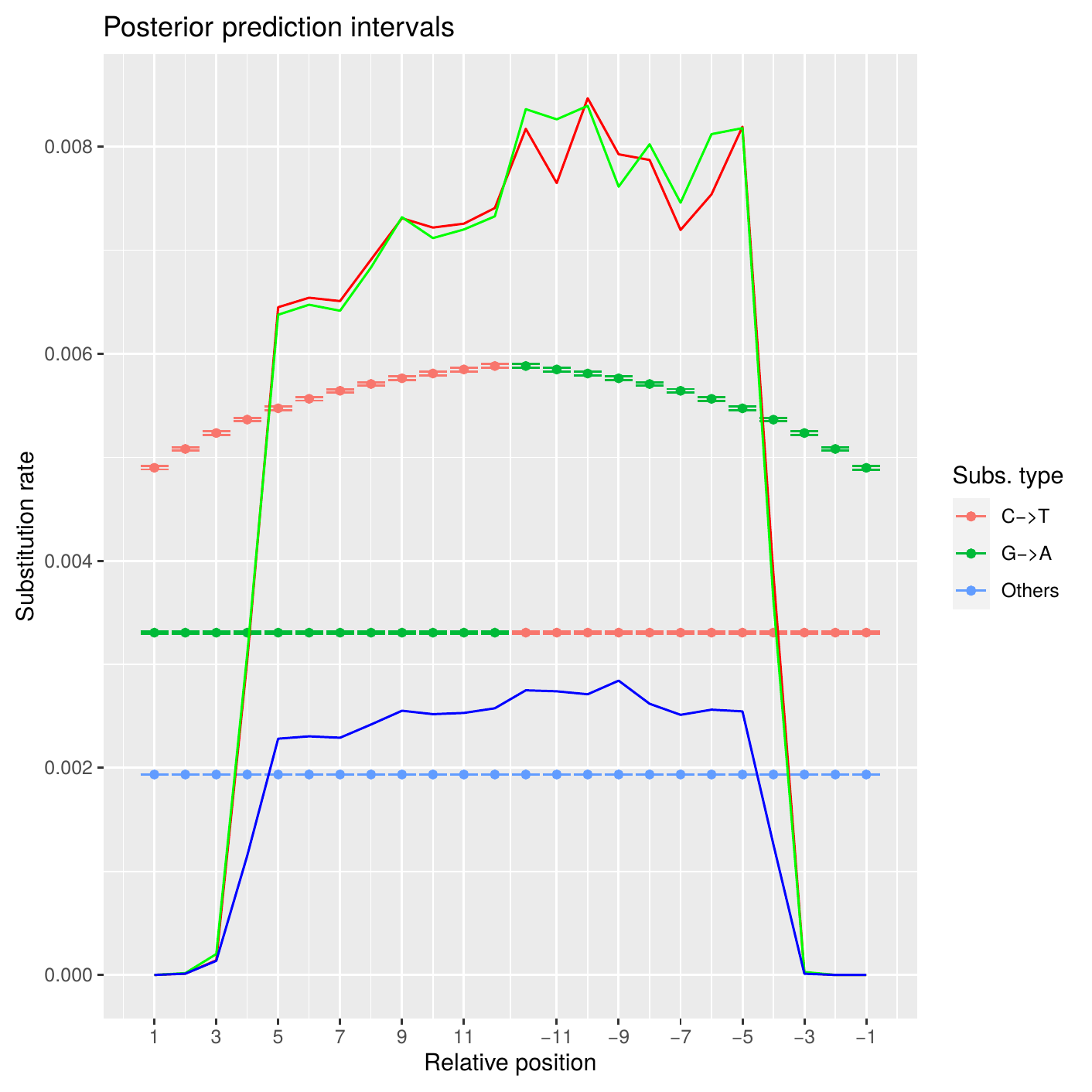

**Figure S6. Posterior predictive check of the mapDamage2.0 damage model for the historical specimen (Rozs3).** Lines and intervals show the observed and predicted per position substitution rates for C to T, G to A and other substitution types as a function of distance from the read ends. Close agreement between observed and predicted values indicates an adequate fit of the damage model.

**Plastome assembly validation and comparison**

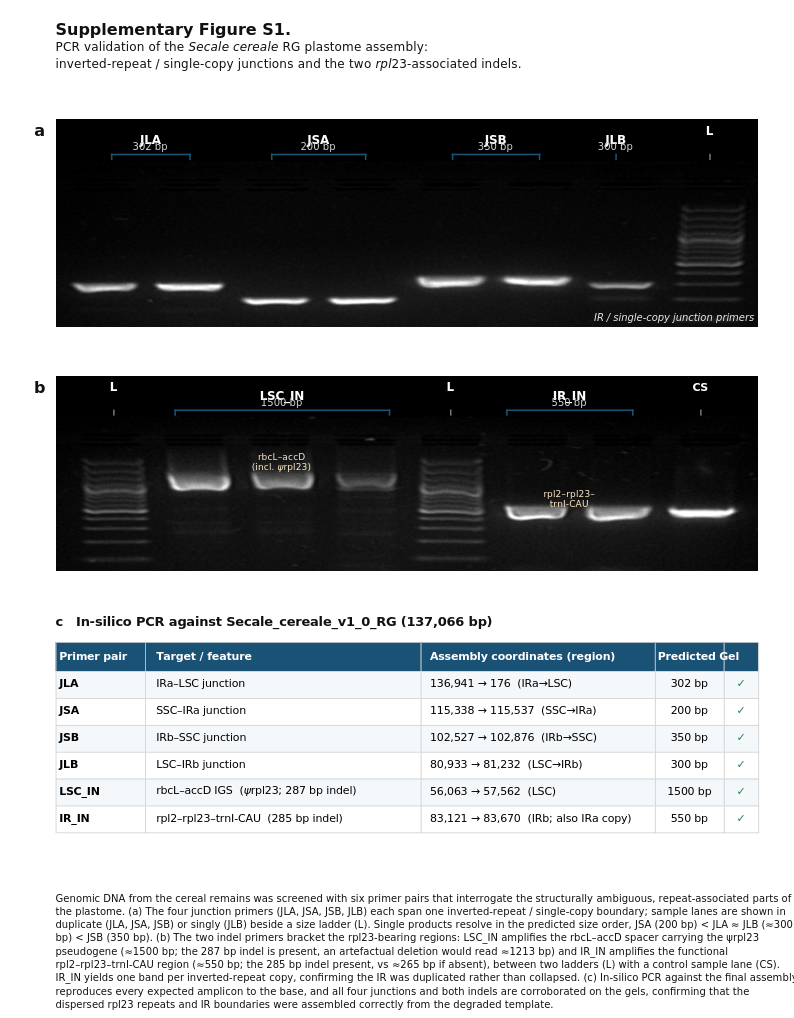

**Figure S7. PCR validation of the Secale cereale RG plastome assembly, inverted-repeat / single-copy junctions and the two rpl23-associated indels.** Genomic DNA from the cereal remains was screened with six primer pairs interrogating the structurally ambiguous, repeat-associated parts of the plastome. (a) The four junction primers (JLA, JSA, JSB, JLB) each span one inverted-repeat / single-copy boundary; sample lanes are shown in duplicate (JLA, JSA, JSB) or singly (JLB) beside a size ladder (L). Single products resolve in the predicted size order, JSA (200 bp) < JLA (about 300 bp) < JSB (350 bp). (b) The two indel primers bracket the *rpl23*-bearing regions: LSC_IN amplifies the *rbcL*–*accD* spacer carrying the ψ*rpl23* pseudogene (about 1500 bp) and IR_IN amplifies the functional *rpl2*–*rpl23*–*trnI*-CAU region (about 550 bp), between two ladders (L) with a control sample lane (CS). (c) In-silico PCR against the final assembly reproduces every expected amplicon, confirming that the dispersed rpl23 repeats and IR boundaries were assembled correctly from the degraded template.

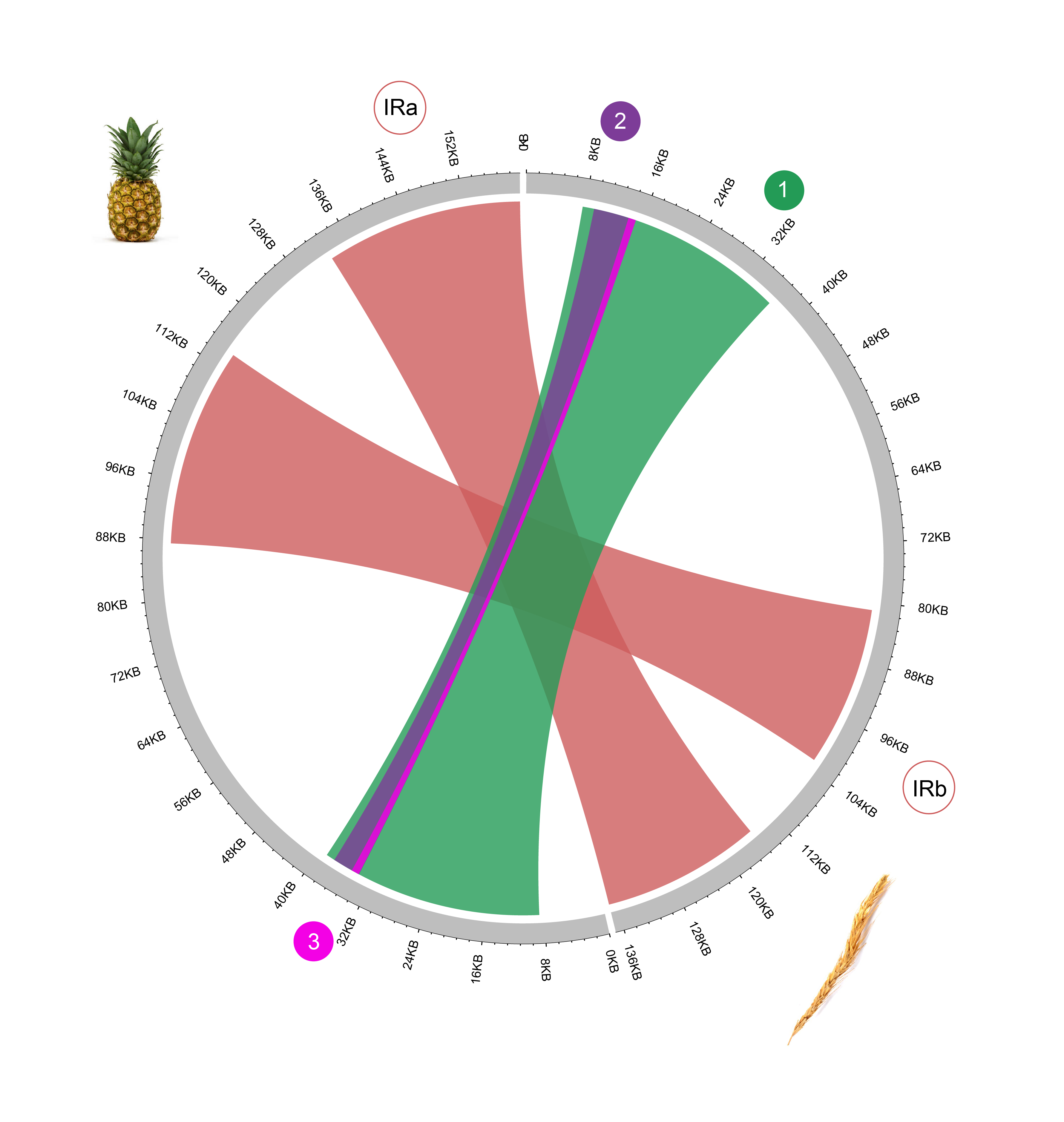

**Figure S8. Collinearity of the historical specimen and Ananas comosus plastid genomes.** The two grey arcs are the plastomes of historical (rye, spike icon) and pineapple (*Ananas comosus*, pineapple icon), scaled in kilobases with the inverted repeats IRa and IRb indicated. Coloured ribbons join homologous blocks; ribbons crossing the centre mark inverted segments. The three large inversions separating grass from bromeliad plastomes (Poczai and Hyvönen 2017) are numbered 1 to 3 (green, purple, magenta), with collinear portions in salmon color.

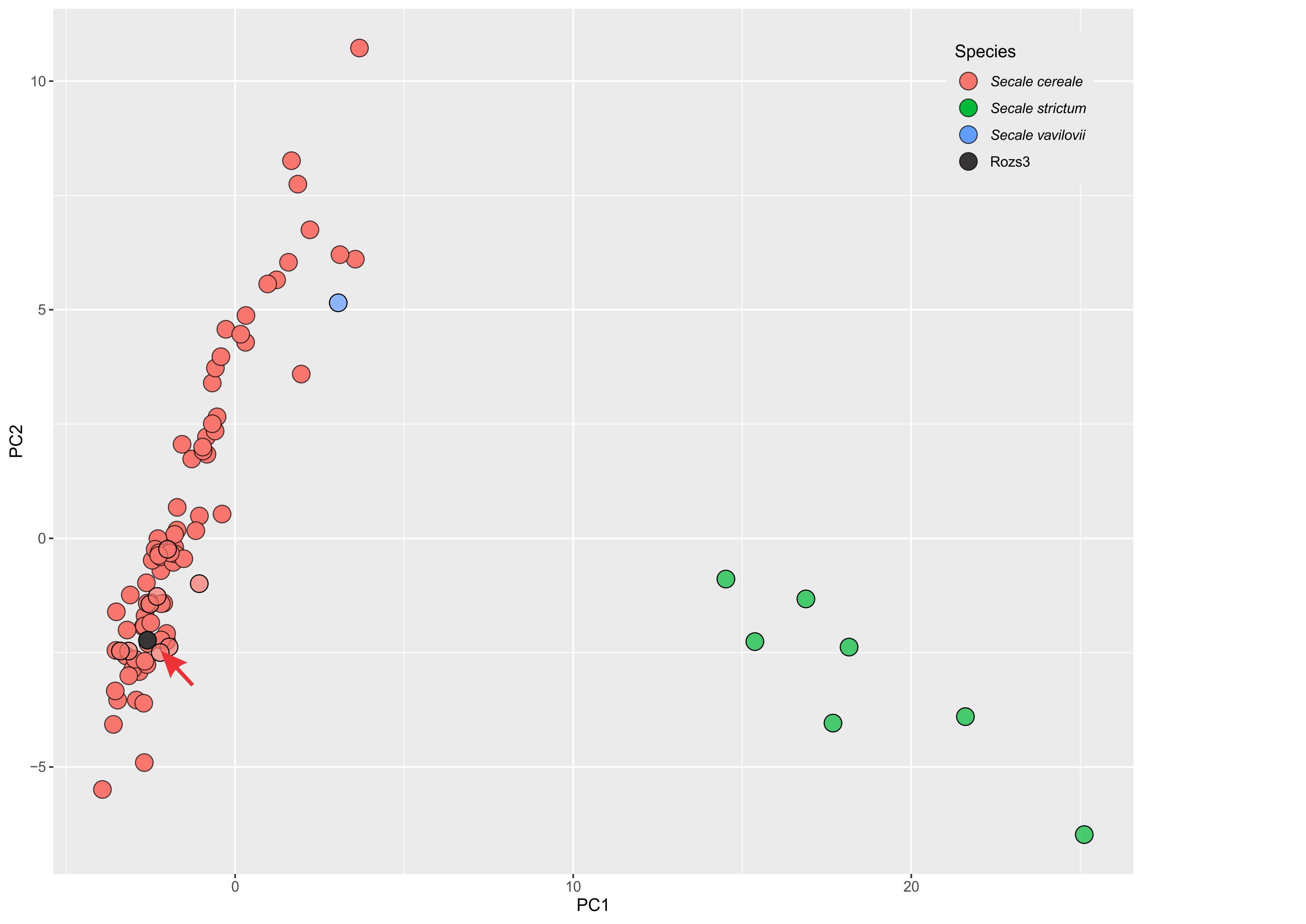

**Figure S9. Species-coloured principal component analysis of 16,198 genome-wide SNPs across the rye panel.** Computed from the same SNP matrix as the country-coloured ordination (Figure 5). *Secale strictum* (green) separates from cultivated *S. cereale* (red) along PC1, with *S. vavilovii* (blue) near the cultivated cloud, so the first component chiefly contrasts wild and cultivated rye. The historical specimen Rozs3 (black, arrowed) falls within the *S. cereale* cluster, consistent with its position in the maximum likelihood tree (Figure 4).

**Supplementary Tables**

**Table S1. Rye accessions used in the genome-wide SNP analysis.** The 92 accessions comprise 84 *Secale cereale* (across the cultivated and weedy subspecies), seven *S. strictum* and one *S. vavilovii*, the last eight serving as outgroups. Accessions are grouped by species and then ordered by country.

| **No.** | **Accession** | **Species** | **Subspecies** | **Country** | **Source panel** | **SRA run** |
| --- | --- | --- | --- | --- | --- | --- |
| 1 | R1038 | *Secale cereale* | subsp. afghanicum | Afghanistan | Schreiber et al. 2019 | ERR2125216 |
| 2 | R1185 | *Secale cereale* | subsp. cereale | Albania | Schreiber et al. 2019 | ERR2125109 |
| 3 | R1213 | *Secale cereale* | subsp. cereale | Albania | Schreiber et al. 2019 | ERR2125560 |
| 4 | R2863 | *Secale cereale* | subsp. dighoricum | Armenia | Schreiber et al. 2019 | ERR2125096 |
| 5 | R875 | *Secale cereale* | subsp. cereale | Austria | Schreiber et al. 2019 | ERR2125183 |
| 6 | R1091 | *Secale cereale* | subsp. cereale | Austria | Schreiber et al. 2019 | ERR2125388 |
| 7 | R34 | *Secale cereale* | subsp. cereale | Canada | Schreiber et al. 2019 | ERR2125574 |
| 8 | R1423 | *Secale cereale* | subsp. cereale | Czech Republic | Schreiber et al. 2019 | ERR2125209 |
| 9 | R1974 | *Secale cereale* | subsp. cereale | Denmark | Rabanus-Wallace et al. (2021) | ERR3663548 |
| 10 | R1341 | *Secale cereale* | subsp. cereale | Estonia | Rabanus-Wallace et al. (2021) | ERR3663379 |
| 11 | R1342 | *Secale cereale* | subsp. cereale | Estonia | Rabanus-Wallace et al. (2021) | ERR3663397 |
| 12 | R864 | *Secale cereale* | subsp. cereale | Georgia | Schreiber et al. 2019 | ERR2125150 |
| 13 | R972 | *Secale cereale* | subsp. cereale | Georgia | Schreiber et al. 2019 | ERR2125516 |
| 14 | R1022 | *Secale cereale* | subsp. cereale | Georgia | Schreiber et al. 2019 | ERR2125375 |
| 15 | R18 | *Secale cereale* | subsp. cereale | Germany | Schreiber et al. 2019 | ERR2125581 |
| 16 | R93 | *Secale cereale* | subsp. cereale | Germany | Schreiber et al. 2019 | ERR2125555 |
| 17 | R100 | *Secale cereale* | subsp. cereale | Germany | Schreiber et al. 2019 | ERR2125080 |
| 18 | R191 | *Secale cereale* | subsp. cereale | Germany | Schreiber et al. 2019 | ERR2125117 |
| 19 | R193 | *Secale cereale* | subsp. cereale | Germany | Schreiber et al. 2019 | ERR2125481 |
| 20 | R1554 | *Secale cereale* | subsp. cereale | Germany | Schreiber et al. 2019 | ERR2125351 |
| 21 | R1572 | *Secale cereale* | subsp. cereale | Germany | Rabanus-Wallace et al. (2021) | ERR3663476 |
| 22 | R1919 | *Secale cereale* | subsp. cereale | Germany | Schreiber et al. 2019 | ERR2125222 |
| 23 | R1968 | *Secale cereale* | subsp. cereale | Germany | Schreiber et al. 2019 | ERR2125311 |
| 24 | R2082 | *Secale cereale* | subsp. cereale | Germany | Schreiber et al. 2019 | ERR2125196 |
| 25 | R2302 | *Secale cereale* | subsp. cereale | Germany | Schreiber et al. 2019 | ERR2125330 |
| 26 | R2413 | *Secale cereale* | subsp. cereale | Germany | Schreiber et al. 2019 | ERR2125404 |
| 27 | R2420 | *Secale cereale* | subsp. cereale | Germany | Schreiber et al. 2019 | ERR2125338 |
| 28 | R2229 | *Secale cereale* | subsp. cereale | Hungary | Schreiber et al. 2019 | ERR2125547 |
| 29 | R2522 | *Secale cereale* | subsp. segetale | Iran | Schreiber et al. 2019 | ERR2125143 |
| 30 | R2581 | *Secale cereale* | subsp. cereale | Iran | Schreiber et al. 2019 | ERR2125163 |
| 31 | R2606 | *Secale cereale* | subsp. ancestrale | Iran | Schreiber et al. 2019 | ERR2125256 |
| 32 | R2743 | *Secale cereale* | subsp. cereale | Iran | Schreiber et al. 2019 | ERR2125169 |
| 33 | R937 | *Secale cereale* | subsp. cereale | Italy | Schreiber et al. 2019 | ERR2125490 |
| 34 | R944 | *Secale cereale* | subsp. cereale | Italy | Schreiber et al. 2019 | ERR2125411 |
| 35 | R1084 | *Secale cereale* | subsp. cereale | Italy | Schreiber et al. 2019 | ERR2125299 |
| 36 | R1090 | *Secale cereale* | subsp. cereale | Italy | Schreiber et al. 2019 | ERR2125138 |
| 37 | R1112 | *Secale cereale* | subsp. cereale | Italy | Schreiber et al. 2019 | ERR2125553 |
| 38 | R1139 | *Secale cereale* | subsp. cereale | Italy | Schreiber et al. 2019 | ERR2125419 |
| 39 | R1069 | *Secale cereale* | subsp. cereale | Lithuania | Rabanus-Wallace et al. (2021) | ERR3663168 |
| 40 | R1304 | *Secale cereale* | subsp. cereale | Lithuania | Schreiber et al. 2019 | ERR3663361 |
| 41 | R1850 | *Secale cereale* | subsp. cereale | Lithuania | Schreiber et al. 2019 | ERR3663530 |
| 42 | R1741 | *Secale cereale* | subsp. cereale | Netherlands | Schreiber et al. 2019 | ERR2125317 |
| 43 | R36 | *Secale cereale* | subsp. cereale | Poland | Schreiber et al. 2019 | ERR2125089 |
| 44 | R37 | *Secale cereale* | subsp. cereale | Poland | Schreiber et al. 2019 | ERR2125496 |
| 45 | R91 | *Secale cereale* | subsp. cereale | Poland | Schreiber et al. 2019 | ERR2125029 |
| 46 | R240 | *Secale cereale* | subsp. cereale | Poland | Schreiber et al. 2019 | ERR2125478 |
| 47 | R242 | *Secale cereale* | subsp. cereale | Poland | Schreiber et al. 2019 | ERR2125130 |
| 48 | R687 | *Secale cereale* | subsp. cereale | Poland | Schreiber et al. 2019 | ERR2125278 |
| 49 | R1105 | *Secale cereale* | subsp. rigidum | Poland | Schreiber et al. 2019 | ERR3663182 |
| 50 | R1319 | *Secale cereale* | subsp. cereale | Poland | Schreiber et al. 2019 | ERR2125540 |
| 51 | R1886 | *Secale cereale* | subsp. cereale | Poland | Schreiber et al. 2019 | ERR2125510 |
| 52 | R1890 | *Secale cereale* | subsp. cereale | Poland | Schreiber et al. 2019 | ERR2125093 |
| 53 | R2190 | *Secale cereale* | subsp. cereale | Poland | Schreiber et al. 2019 | ERR2125391 |
| 54 | R2203 | *Secale cereale* | subsp. cereale | Poland | Schreiber et al. 2019 | ERR2125324 |
| 55 | R1133 | *Secale cereale* | subsp. cereale | Portugal | Schreiber et al. 2019 | ERR2125565 |
| 56 | R2275 | *Secale cereale* | subsp. cereale | Portugal | Schreiber et al. 2019 | ERR2125371 |
| 57 | R684 | *Secale cereale* | subsp. cereale | Romania | Schreiber et al. 2019 | ERR2125271 |
| 58 | R1201 | *Secale cereale* | subsp. cereale | Romania | Schreiber et al. 2019 | ERR2125431 |
| 59 | R20 | *Secale cereale* | subsp. cereale | Russia | Schreiber et al. 2019 | ERR2125594 |
| 60 | R31 | *Secale cereale* | subsp. cereale | Russia | Schreiber et al. 2019 | ERR2125022 |
| 61 | R47 | *Secale cereale* | subsp. cereale | Russia | Schreiber et al. 2019 | ERR2125047 |
| 62 | R233 | *Secale cereale* | subsp. cereale | Russia | Schreiber et al. 2019 | ERR2125503 |
| 63 | R235 | *Secale cereale* | subsp. cereale | Russia | Schreiber et al. 2019 | ERR2125102 |
| 64 | R1248 | *Secale cereale* | subsp. cereale | Russia | Schreiber et al. 2019 | ERR2125176 |
| 65 | R1257 | *Secale cereale* | subsp. cereale | Russia | Schreiber et al. 2019 | ERR2125202 |
| 66 | R1384 | *Secale cereale* | subsp. cereale | Russia | Schreiber et al. 2019 | ERR2125554 |
| 67 | R1450 | *Secale cereale* | subsp. cereale | Russia | Schreiber et al. 2019 | ERR2125288 |
| 68 | R1452 | *Secale cereale* | subsp. cereale | Russia | Schreiber et al. 2019 | ERR2125568 |
| 69 | R1972 | *Secale cereale* | subsp. cereale | Russia | Schreiber et al. 2019 | ERR2125358 |
| 70 | R1987 | *Secale cereale* | subsp. cereale | Russia | Schreiber et al. 2019 | ERR2125297 |
| 71 | R2136 | *Secale cereale* | subsp. cereale | Russia | Schreiber et al. 2019 | ERR2125236 |
| 72 | R604 | *Secale cereale* | subsp. cereale | Slovakia | Schreiber et al. 2019 | ERR2125304 |
| 73 | R607 | *Secale cereale* | subsp. segetale | Slovakia | Schreiber et al. 2019 | ERR2125384 |
| 74 | R612 | *Secale cereale* | subsp. cereale | Slovakia | Schreiber et al. 2019 | ERR2125345 |
| 75 | R831 | *Secale cereale* | subsp. cereale | Slovakia | Schreiber et al. 2019 | ERR2125136 |
| 76 | R779 | *Secale cereale* | subsp. segetale | Spain | Schreiber et al. 2019 | ERR2125364 |
| 77 | R783 | *Secale cereale* | subsp. cereale | Spain | Schreiber et al. 2019 | ERR2125284 |
| 78 | R788 | *Secale cereale* | subsp. segetale | Spain | Schreiber et al. 2019 | ERR2125250 |
| 79 | R2449 | *Secale cereale* | subsp. cereale | Spain | Schreiber et al. 2019 | ERR2125189 |
| 80 | R1001 | *Secale cereale* | subsp. cereale | Switzerland | Schreiber et al. 2019 | ERR2125123 |
| 81 | R253 | *Secale cereale* | subsp. cereale | Turkey | Schreiber et al. 2019 | ERR2125010 |
| 82 | R280 | *Secale cereale* | subsp. cereale | Turkey | Schreiber et al. 2019 | ERR2125588 |
| 83 | R1148 | *Secale cereale* | subsp. cereale | Turkey | Schreiber et al. 2019 | ERR2125441 |
| 84 | R1429 | *Secale cereale* | subsp. cereale | Ukraine | Schreiber et al. 2019 | ERR2125477 |
| 85 | R579 | *Secale strictum* | subsp. kuprijanovii | Azerbaijan | Rabanus-Wallace et al. (2021) | ERR3663872 |
| 86 | R2431 | *Secale strictum* | subsp. strictum | Bulgaria | Rabanus-Wallace et al. (2021) | ERR3663656 |
| 87 | R797 | *Secale strictum* | subsp. anatolicum | Poland | Rabanus-Wallace et al. (2021) | ERR3664016 |
| 88 | R1053 | *Secale strictum* | subsp. kuprijanovii | Slovenia | Rabanus-Wallace et al. (2021) | ERR3663116 |
| 89 | R264 | *Secale strictum* | subsp. africanum | South Africa | Rabanus-Wallace et al. (2021) | ERR3663746 |
| 90 | R1210 | *Secale strictum* | subsp. africanum | South Africa | Rabanus-Wallace et al. (2021) | ERR3663336 |
| 91 | R1119 | *Secale strictum* | subsp. anatolicum | Turkey | Rabanus-Wallace et al. (2021) | ERR3663230 |
| 92 | R1063 | *Secale vavilovii* | — | Poland | Rabanus-Wallace et al. (2021) | ERR3663135 |

**Table S2. Polymorphism counts for each of the 36 selective sweep regions.** Regions identified in the Weining rye genome (Li et al. 2021), named by Weining gene identifier, chromosome and detection statistic (DRI, XP or FST). Length is the region size in base pairs. All values are numbers of polymorphic sites measured relative to the historical specimen (Rozs3): the Weining column gives sites where the historical rye differs from the Weining reference, and the R2446, R925 and PUMA columns sites where each modern cultivar differs from the reference-guided consensus of the historical rye. Cultivar reads were obtained from Rabanus-Wallace et al. (2021), accessions R2446 (ERR3771531), R925 (ERR3771534) and PUMA (SRR10088797).

| **Identified selected region** | **Length (bp)** | **Chr** | **Class** | **Weining** | **R2446** | **R925** | **PUMA** |
| --- | --- | --- | --- | --- | --- | --- | --- |
| ScWN1R01G158700_Chr1_DRI | 2,742 | 1 | DRI | 13 | 35 | 18 | 9 |
| ScWN1R01G240500_Chr1_DRI | 25,082 | 1 | DRI | 724 | 403 | 343 | 131 |
| ScWN1R01G314500_Chr1_DRI | 3,765 | 1 | DRI | 18 | 26 | 22 | 1 |
| ScWN1R01G420200_Chr1_DRI | 1,753 | 1 | DRI | 69 | 66 | 38 | 67 |
| ScWN1R01G474300_Chr1_XP | 3,434 | 1 | XP | 50 | 76 | 61 | 30 |
| ScWN1R01G523000_Chr1_XP | 13,431 | 1 | XP | 52 | 320 | 635 | 107 |
| ScWN2R01G087200_Chr2_XP | 1,524 | 2 | XP | 16 | 67 | 22 | 10 |
| ScWN2R01G169300_Chr2_XP | 12,688 | 2 | XP | 435 | 233 | 290 | 161 |
| ScWN2R01G177000_Chr2_DRI | 3,339 | 2 | DRI | 24 | 101 | 68 | 15 |
| ScWN2R01G187200_Chr2_FST | 2,792 | 2 | FST | 27 | 34 | 21 | 11 |
| ScWN2R01G269300_Chr2_DRI | 1,191 | 2 | DRI | 0 | 9 | 5 | 0 |
| ScWN2R01G313400_Chr2_FST | 4,146 | 2 | FST | 41 | 72 | 26 | 2 |
| ScWN2R01G422000_Chr2_XP | 4,310 | 2 | XP | 100 | 120 | 107 | 69 |
| ScWN3R01G102300_Chr3_XP | 1,170 | 3 | XP | 18 | 47 | 13 | 8 |
| ScWN3R01G147500_Chr3_XP | 808 | 3 | XP | 0 | 3 | 3 | 0 |
| ScWN3R01G332700_Chr3_FST | 4,010 | 3 | FST | 48 | 123 | 118 | 32 |
| ScWN3R01G377900_Chr3_XP | 11,386 | 3 | XP | 121 | 292 | 171 | 118 |
| ScWN3R01G411500_Chr3_XP | 738 | 3 | XP | 2 | 1 | 4 | 0 |
| ScWN4R01G046700_Chr4_XP | 5,311 | 4 | XP | 117 | 197 | 136 | 64 |
| ScWN4R01G066200_Chr4_XP | 7,570 | 4 | XP | 32 | 85 | 38 | 11 |
| ScWN4R01G072600_Chr4_XP | 3,334 | 4 | XP | 60 | 77 | 81 | 25 |
| ScWN4R01G247400_Chr4_XP | 4,850 | 4 | XP | 48 | 101 | 68 | 25 |
| ScWN4R01G449800_Chr4_XP | 5,080 | 4 | XP | 140 | 215 | 132 | 68 |
| ScWN5R01G089500_Chr5_DRI | 474 | 5 | DRI | 2 | 4 | 4 | 1 |
| ScWN5R01G132700_Chr5_DRI | 1,664 | 5 | DRI | 7 | 14 | 11 | 2 |
| ScWN5R01G313900_Chr5_XP | 3,532 | 5 | XP | 47 | 76 | 111 | 23 |
| ScWN5R01G347500_Chr5_XP | 1,806 | 5 | XP | 27 | 31 | 26 | 18 |
| ScWN6R01G057200_Chr6_DRI | 2,414 | 6 | DRI | 38 | 49 | 31 | 37 |
| ScWN6R01G089100_Chr6_DRI | 9,605 | 6 | DRI | 20 | 25 | 16 | 1 |
| ScWN6R01G115700_Chr6_DRI | 54,574 | 6 | DRI | 855 | 494 | 239 | 90 |
| ScWN6R01G233700_Chr6_DRI | 6,331 | 6 | DRI | 8 | 30 | 25 | 15 |
| ScWN6R01G470400_Chr6_XP | 1,142 | 6 | XP | 26 | 33 | 14 | 9 |
| ScWN7R01G256000_Chr7_DRI | 1,814 | 7 | DRI | 2 | 9 | 3 | 1 |
| ScWN7R01G263700_Chr7_FST | 3,128 | 7 | FST | 11 | 14 | 43 | 0 |
| ScWN7R01G370700_Chr7_FST | 6,210 | 7 | FST | 10 | 54 | 17 | 7 |
| ScWN7R01G441300_Chr7_XP | 2,842 | 7 | XP | 23 | 90 | 92 | 19 |

**Table S3. Polymorphism counts and densities summarised by chromosome.** For each chromosome, the combined length of its sweep regions and, for every line, the total polymorphic sites relative to the historical rye with density as a percentage of region length. Counts and comparisons as in Table S2.

| **Chromosome** | **Size (bp)** | **Weining** | **%** | **R2446** | **%** | **R925** | **%** | **PUMA** | **%** |
| --- | --- | --- | --- | --- | --- | --- | --- | --- | --- |
| Chromosome 1 | 50,207 | 926 | 1.84 | 936 | 1.86 | 1,117 | 2.22 | 345 | 0.68 |
| Chromosome 2 | 29,990 | 643 | 2.14 | 636 | 2.12 | 539 | 1.79 | 268 | 0.89 |
| Chromosome 3 | 17,059 | 189 | 1.10 | 466 | 2.73 | 309 | 1.81 | 158 | 0.92 |
| Chromosome 4 | 26,145 | 397 | 1.51 | 675 | 2.54 | 455 | 1.74 | 193 | 0.73 |
| Chromosome 5 | 7,476 | 83 | 1.11 | 125 | 1.67 | 152 | 2.03 | 44 | 0.58 |
| Chromosome 6 | 74,066 | 947 | 1.27 | 631 | 0.85 | 325 | 0.43 | 152 | 0.20 |
| Chromosome 7 | 13,994 | 46 | 0.32 | 167 | 1.19 | 155 | 1.10 | 27 | 0.19 |

**Table S4. Polymorphism counts and densities summarised by sweep class.** The 36 regions grouped by detection statistic (13 DRI, 18 XP, 5 FST). For each class, the number of regions, combined length, and per line the total polymorphic sites relative to the historical rye with density as a percentage of class length. The final row pools all regions. Counts and comparisons as in Table S2.

| **Sweep class** | **Regions (n)** | **Size (bp)** | **Weining** | **%** | **R2446** | **%** | **R925** | **%** | **PUMA** | **%** |
| --- | --- | --- | --- | --- | --- | --- | --- | --- | --- | --- |
| DRI | 13 | 114,748 | 1,780 | 1.55 | 1,265 | 1.10 | 823 | 0.72 | 370 | 0.32 |
| XP | 18 | 84,956 | 1,314 | 1.55 | 2,064 | 2.43 | 2,004 | 2.36 | 765 | 0.90 |
| FST | 5 | 20,286 | 137 | 0.68 | 297 | 1.46 | 225 | 1.11 | 52 | 0.26 |
| All regions | 36 | 219,990 | 3,231 | 1.47 | 3,626 | 1.65 | 3,052 | 1.39 | 1,187 | 0.54 |
